# Leukemia core transcriptional circuitry is a sparsely interconnected hierarchy stabilized by incoherent feed-forward loops

**DOI:** 10.1101/2023.03.13.532438

**Authors:** Taku Harada, Jérémie Kalfon, Monika W. Perez, Kenneth Eagle, Flora Dievenich Braes, Rashad Batley, Yaser Heshmati, Juliana Xavier Ferrucio, Jazmin Ewers, Stuti Mehta, Andrew Kossenkov, Jana M. Ellegast, Allyson Bowker, Jayamanna Wickramasinghe, Behnam Nabet, Vikram R. Paralkar, Neekesh V. Dharia, Kimberly Stegmaier, Stuart H. Orkin, Maxim Pimkin

## Abstract

Lineage-defining transcription factors form densely interconnected circuits in chromatin occupancy assays, but the functional significance of these networks remains underexplored. We reconstructed the functional topology of a leukemia cell transcription network from the direct gene-regulatory programs of eight core transcriptional regulators established in pre-steady state assays coupling targeted protein degradation with nascent transcriptomics. The core regulators displayed narrow, largely non-overlapping direct transcriptional programs, forming a sparsely interconnected functional hierarchy stabilized by incoherent feed-forward loops. BET bromodomain and CDK7 inhibitors disrupted the core regulators’ direct programs, acting as mixed agonists/antagonists. The network is predictive of dynamic gene expression behaviors in time-resolved assays and clinically relevant pathway activity in patient populations.

## Introduction

Cell type-specific gene expression programs in metazoans are established by relatively small sets of lineage-restricted transcription factors (TFs), variably referred to as “core regulatory”, “master” or “reprogramming” (*1*–*8*).How TFs cooperate to form a gene regulatory network is fundamental to understanding development and disease (*9*–*11*). TFs directly interpret the genome by binding to cognate DNA sequences (TF motifs) in genomic regulatory elements to activate or repress transcription of nearby genes (*12*). Although many studies have functionally linked individual TF binding events to changes in gene expression (*13*–*15*), establishing the full complement of a TF’s direct genomic targets remains challenging. Genetic knockouts and forced expression systems have been used to probe the overall functional significance of TFs, but the slow kinetics of these assays precludes distinguishing direct TF targets from secondary effects (*16*). In practice, binding of a TF to a gene’s promoter or enhancer, whether observed in a chromatin occupancy assay or inferred from the presence of its cognate DNA motif, has been commonly interpreted as evidence of direct regulation. From this perspective, genome-scale chromatin occupancy maps have been used to reconstruct gene regulatory networks, revealing a common topology shared between various cell types and organisms: a small, densely interconnected core of master TFs making cooperative connections to an extended network of peripheral genomic targets (*5*, *17*–*21*). Based on these observations, a pervasive model posits that core TFs directly enforce their own and each other’s expression, forming a self-sustaining network of interconnected feedback and feed-forward loops termed core regulatory circuitry (CRC) (*5*, *22*–*28*). However, as CRCs are ultimately inferred from TF binding patterns, the extent to which they include functional regulatory connections has not been rigorously tested. Meanwhile, emerging data indicate that TFs may directly regulate only a small fraction of the genes they occupy (*29*–*31*), suggesting that the true topology of a transcription network would be more accurately constructed from functionally established direct programs of each regulator.

We sought to systematically deconvolute a transcription regulatory network by elucidating the direct programs of multiple core transcriptional regulators (CTRs) in a single human cell context. Using acute myeloid leukemia (AML) as a model system, we engineered chemical degrons of eight CTRs and established their direct gene-regulatory functions in pre-steady state assays coupling rapid protein degradation with nascent transcriptomics. Rather than the canonical CRC model, our data reveal a sparsely interconnected hierarchy of core regulators characterized by the presence of both positive and negative regulatory connections, lack of positive self-regulation, and much narrower direct transcriptional programs than previously inferred.

## Results

### Definition of an AML core regulatory circuitry

We began by defining a network of core regulatory TFs fulfilling the most stringent canonical CRC criteria: *1*) association of coding genes with extended enhancers characterized by disproportionately strong histone acetylation and cofactor recruitment, termed superenhancers or stretch enhancers, *2*) binding to their own and each other’s superenhancers and *3*) lineage-selective dependency (*22*, *23*). In recent work, we integrated gene dependency and chromatin architecture datasets to define a putative 29-member CRC in AML (*31*). Briefly, among a chromatin immunoprecipitation sequencing (ChIP-seq) data set for the enhancer histone mark H3K27ac in a panel of 130 primary AML samples, PDX models and cell lines, we identified 561 TF-coding genes associated with recurrent AML superenhancers. We then intersected the 561 superenhancer-associated TF-coding genes with genes identified to be selective dependencies for AML versus other malignancies in the Broad Cancer Dependency Map project, resulting in a list of 29 putative core TFs (Figs. 1A, S1) (*31*). We further hypothesized that lineage-restricted transcriptional cofactors may have highly specialized regulatory roles similar to those of lineage TFs and extended the list by including 3 non-TF cofactors fulfilling the above criteria (IRF2BP2, LMO2 and ZMYND8). Further including the pan-essential critical oncogenic TFs MYC and MAX resulted in a 34-member extended AML CRC (Fig. S2). Because this putative circuit included transcriptional cofactors in addition to DNA-binding TFs, we adopted the term “core transcriptional regulator” (CTR). We examined the binding preferences of 27 of the 34 CTRs by ChIP-seq in an AML cell line (MV411), detecting dense co-occupancy of CTRs in DNA elements (Figs. S3-S4). Fourteen CTRs, including 12 TFs and 2 cofactors, formed a fully interconnected circuitry of feedback and feed-forward superenhancer occupancy loops, thus constituting a minimal CRC fulfilling the canonical criteria in MV411 cells (Fig. 1B,C).

**Figure 1.**
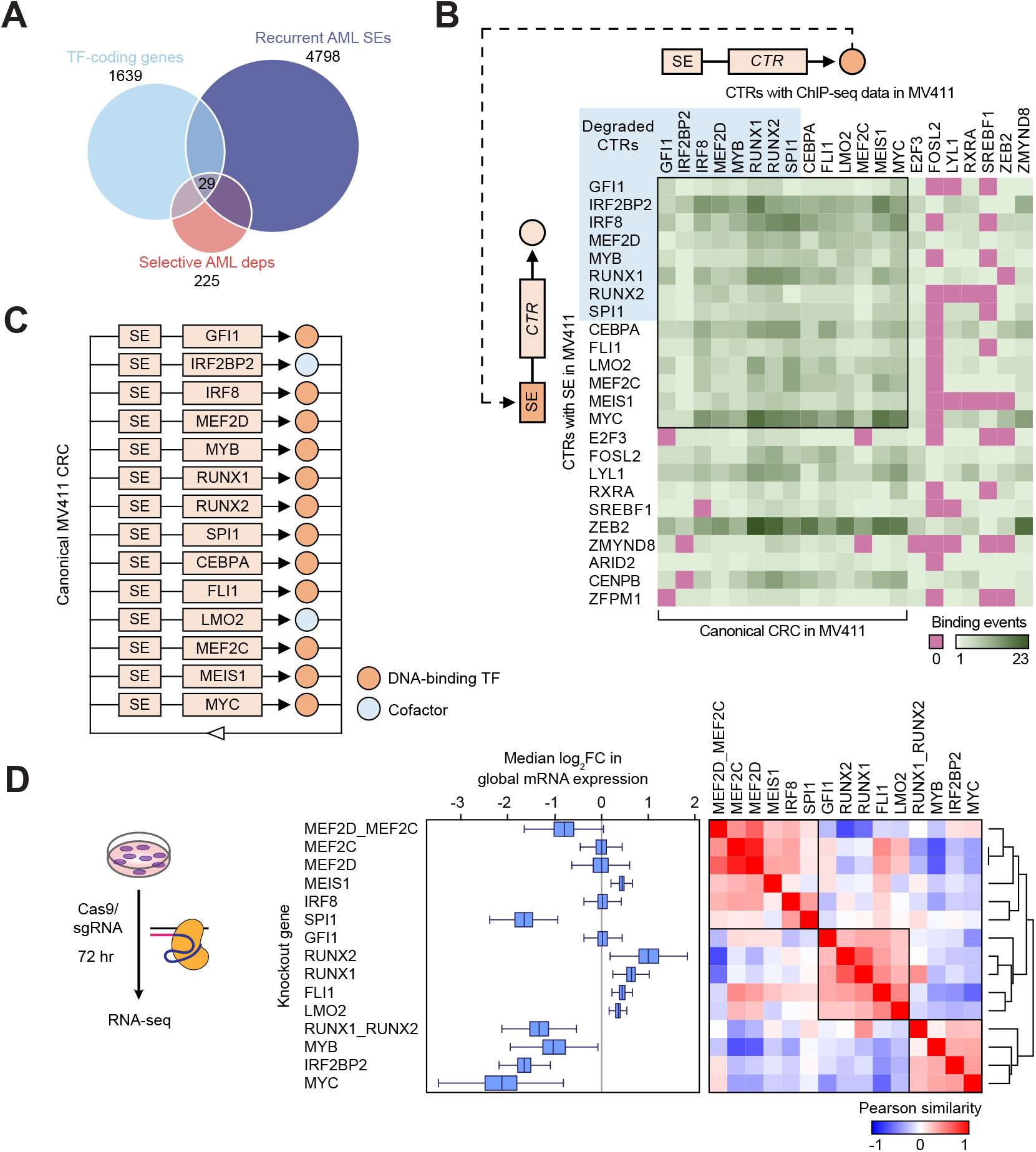
A canonical AML CRC. A, Definition of candidate core TFs at the intersection of recurrent AML superenhancers, enriched AML dependencies and human TF-coding genes. B, Connectivity heatmap visualizing TFs (columns) binding to their own and each other’s superenhancers (rows). Individual cells visualize the number of binding events detected by ChIP-seq in MV411 cells (Methods). CTR genes not associated with superenhancers in MV411 cells were excluded from the analysis. C, A typical visualization of a minimal AML CRC fulfilling the canonical criteria, illustrating complete interconnectivity between the core members. D, MV411 cells were electroporated with *in vitro* assembled Cas9/sgRNA complexes targeting individual CTRs and combinations of TF paralogs and RNA-seq was performed 72 hours after electroporation. Validation of knockout efficiency by Western blot is found in Fig. S5. The boxplots visualize knockout-induced changes in the expression of top 5000 expressed genes measured by RNA-seq normalized to an external spike-in control. The elements of the boxplots are as follows: center line, median log_2_ fold change in mRNA expression; box limits, upper and lower quartiles; whiskers, 1.5x interquartile range. The heatmap visualizes TF knockouts hierarchically clustered by Pearson correlation between knockout-induced changes in the expression of the top 5000 expressed genes.

### CRC displays functional modularity and redundancy

We examined the long-term functional consequences of CTR depletion by inactivating 13 of the 14 canonical CTRs by CRISPR/Cas9 editing and measuring the mRNA pools by RNA-seq (Fig. 1D, S5). To evaluate the degree of global transcriptional response we normalized transcript counts to external spike-in controls (*32*, *33*). Knockouts of MYB, MYC, SPI1 and IRF2BP2 resulted in a transcriptome-wide collapse. Concurrent knockouts of TF paralogs (MEF2D/MEF2C and RUNX1/RUNX2) led to a similar outcome, consistent with their synthetic lethality and indicating partially redundant functions (Fig. 1D, S5). Similar trends were observed on quantitative measurements of genome-wide histone acetylation (Fig. S5). To assess the degree of functional divergence within the CRC, we asked whether individual regulators could be grouped into functional modules based on the patterns of co-regulated effector genes. Indeed, unsupervised clustering of knockout-induced mRNA changes revealed a modular structure of regulator-target relationships, where the CTRs formed three partially antagonistic modules (Fig. 1D). Thus, although the steady-state transcriptional responses to genetic perturbation provided no information on the CTRs’ direct transcriptional programs, they uncovered prominent functional divergence and modularity within the CRC, pointing to a more complex core regulatory structure than implied by the canonical “one for all and all for one” model.

### CTRs have narrow direct transcriptional programs

To define the direct gene-regulatory functions of individual CTRs we utilized direct measurements of genome-wide transcription rates following rapid targeted protein degradation (*29*). Using MV411 as a parent line, we engineered eight model cell lines, each with a CTR (MYB, IRF2BP2, SPI1/PU.1, RUNX1, RUNX2, IRF8, MEF2D or GFI1) fused to an FKBP12^F36V^ (dTAG) domain and a fluorescent tag by a homozygous knock-in of the FKBP12^F36V^-mScarlet-coding DNA sequence into the endogenous CTR locus (Fig. 2A, S6). Treatment with dTAG^V^-1, a VHL-engaging PROTAC (*34*), led to a near-complete loss of the fusion proteins within 1-3 hours (Fig 2B, S6). Following rapid degradation of CTRs, we measured genome-wide rates of mRNA synthesis by SLAM-seq (*35*). We defined direct genomic targets of each CTR as those genes which displayed significant changes in transcription rates (FDR<0.05) immediately after a near-complete CTR degradation. Strikingly, all eight CTRs displayed narrow direct gene-regulatory programs, affecting 15 to 450 target genes (Fig. 2C). Each CTR displayed evidence of both direct gene repression and activation, including CTRs generally considered to be transcriptional repressors (IRF2BP2, GFI1) and activators (MYB). The 8 CTRs collectively regulated a total of 1076 target genes, or 14% of the expressed genome. The direct transcriptional programs of individual CTRs were largely non-overlapping: of the 1076 genes regulated by at least 1 CTR, only 274 genes were affected by 2 or more CTRs, resulting in a very low-connectivity network (1510 edges for 1079 nodes, or 1.4 edges/node; Fig. 3D-G). When two CTRs converged on a small set of overlapping targets, they generally displayed a mix of concordant and discordant behaviors (Fig. 2H). The RUNX paralogs displayed largely distinct direct transcriptional programs, but highly concordant behavior at overlapping targets (Fig. 2G,H). The functional network, although more interconnected than expected by chance (Fig. 2E), contrasts with the dramatically larger and denser network formed by the same 8 CTRs in chromatin binding assays (52960 edges for 7130 nodes, corresponding to 95% of the expressed genome at an average connectivity of 7.4 edges/node). We conclude that the AML CTRs display universally narrow and largely distinct direct transcriptional programs. In agreement with prior reports (*29*, *31*), chromatin occupancy by itself appears to be a poor predictor of direct regulation.

**Figure 2.**
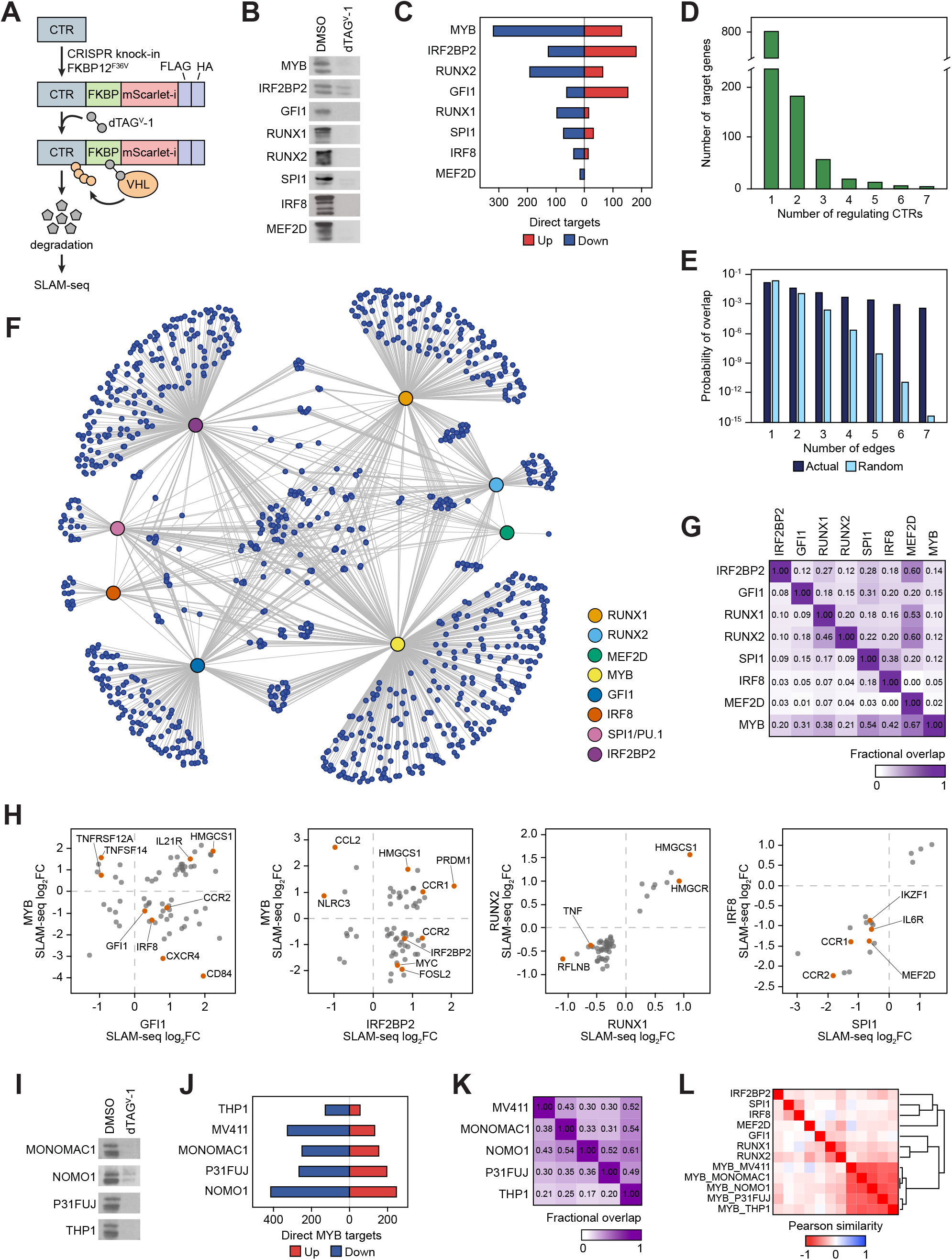
A direct gene-regulatory network in AML. A, Schematic of endogenous CTR degron tagging by CRISPR-HDR and subsequent targeted degradation of the fusion proteins. Refer to Fig. S6 for details of degron validation. B, Western blot demonstrating degradation of CTR-FKBP fusions by dTAG^v^-1. The time points, chosen as the earliest point at which a near-complete degradation of a CTR is observed, were as follows: 1 hr: MYB, SPI1, RUNX1, RUNX2; 2 hrs: GFI1, IRF8, MEF2D; 3 hrs: IRF2BP2. C, Numbers of direct targets of the 8 CTRs visualized according to the direction of response to CTR degradation. Downregulated targets represent genes activated by the CTR, whereas upregulated targets represent genes repressed by the CTR. D, Overlap between the direct targets of 8 CTRs, visualized as number of genes directly regulated by *n* CTRs. E, Actual vs. randomly expected overlap between the direct targets of 8 CTRs, visualized as fraction of the expressed genome co-regulated by *n* CTRs. F, A visualization of the reconstructed core regulatory network, where large nodes represent CTRs, small nodes represent peripheral targets and edges represent direct regulatory connections. G, Fractional pairwise overlap between the direct transcriptional programs of the 8 CTRs. Data points reflect fraction of columns overlapping with rows. H, Nascent transcriptomics responses of overlapping direct targets to CTR degradation, demonstrating mixed effects of the CTRs on co-regulated genes. Data points reflect log_2_ fold change in gene transcription rates immediately after CTR degradation by dTAG^v^-1 compared to vehicle control. I, Western blot demonstrating degradation of MYB-dTAG fusions by dTAG^v^-1 in 4 additional cell lines. J, Direct MYB targets in 5 cell lines are visualized according to the direction of response to MYB degradation. K, Fractional pairwise overlap between the direct transcriptional programs of MYB in 5 cell line models. Data points reflect fraction of columns overlapping with rows. L, Pearson similarity of the genome-scale changes in nascent transcription rates following CTR degradation.

**Figure 3.**
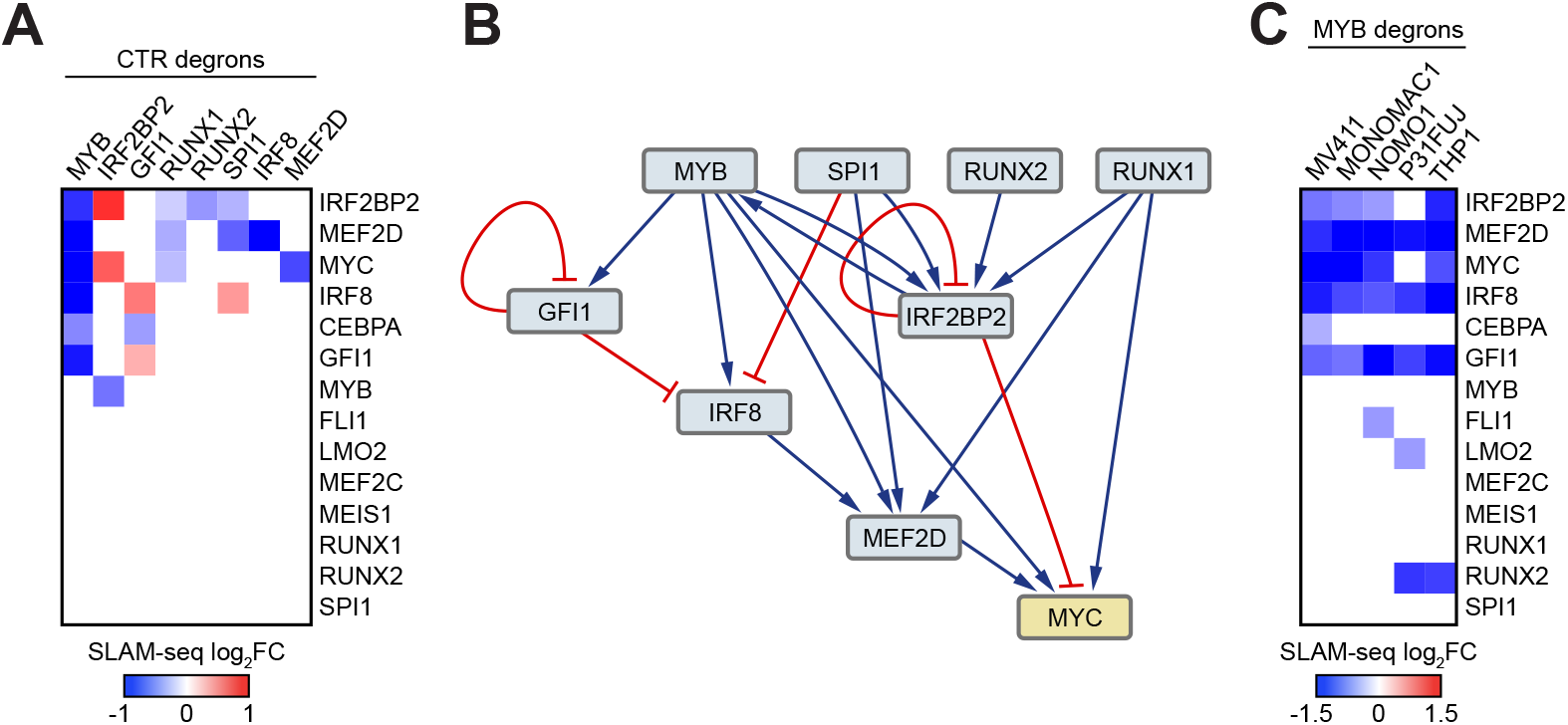
Functional CRC topology. A, Transcription rate response of the 14 core members to CTR degradation. The heatmap visualizes log_2_ fold change in nascent transcript counts immediately after CTR degradation by dTAG^v^-1. Only significant (BH-adjusted *p*-value <0.05) changes are shown. B, Visualization of the direct regulatory relationships between the 8 degraded CTRs. Blue arrows represent positive (activating) connections while red connectors represent inhibiting (repressive) connections. C, Transcription rate response of the 14 core members to MYB degradation in 5 AML cell line models. The heatmap visualizes log_2_ fold change in nascent transcript counts after a 1-hour MYB degradation. Only significant (BH-adjusted *p*-value <0.05) changes are shown.

We sought to evaluate the degree to which the direct transcriptional program of a CTR may be context dependent. We engineered MYB degrons and established direct MYB targets in 4 additional AML cell lines (Figs. 2I,J, S7). While the pairwise overlap between direct MYB targets in the 5 cell lines ranged between 17-61% (Fig. 2K, S7), the definition of a direct target depends on the significance cutoff, and a simple overlap likely underestimates the functional similarities between MYB programs. Indeed, the 5 MYB degrons displayed strong similarity on pairwise correlation of genome-wide transcription rates (Fig. 2L). This observation indicates that the MYB program is largely hard-wired, but the amplitude of MYB effect at the level of individual genes is modulated by the cell context.

### A sparsely interconnected, hierarchical CRC structure

We sought next to reconstruct the core regulatory structure by examining how each CTR regulated transcription of the other core members. In contrast to the occupancy-based CRC model stipulating that each core regulator directly enforces its own expression and the expression of all other core members, each of the 8 degraded CTRs directly impacted only between 1 and 6 other members of the 14-member CRC (Fig. 3A). Furthermore, contrary to the canonical model, we detected both positive (activating) and negative (repressive) regulatory connections. Only two CTRs, IRF2BP2 and GFI1, displayed evidence of self-regulation; both inhibited, rather than activated, their own transcription. The CRC displayed a cascading hierarchical structure: three TFs (RUNX1, RUNX2 and SPI1) showed no evidence of regulation by another CTR, whereas IRF2BP2, MEF2D and MYC were each a point of convergence of 4 regulating CTRs (Fig. 3B). IRF2BP2 and MYB formed a positive reciprocal feedback loop, representing the only example of direct reciprocal regulation we found. Notably, the network characteristics of the transcriptional cofactor IRF2BP2 were indistinguishable from the DNA-binding core TFs. Indeed, IRF2BP2 demonstrated a similarly narrow direct transcriptional program and appeared to be an integral member of the CRC, participating in both gene activation and repression. While the overall degree of connectivity was higher between the CRC members compared to the extended CRC network (2.1 edges/node vs. 1.4 edges/node), it was still markedly lower than would be expected in a fully interconnected circuit (9 edges/node). Consistent with the genome-wide trends, the direct connections between MYB and the other core members were largely conserved between the 5 MYB degron cell models (Fig. 3C). Collectively, these observations paint a fundamentally different picture of the functional CRC topology than previously inferred principally from TF occupancy data.

### CRC is stabilized by incoherent feed-forward loops

So far, our data indicate that prolonged depletion of some CTRs leads to a global transcriptional collapse while rapid-kinetics experiments demonstrate that CTRs have universally narrow direct transcriptional programs. These observations indicate that the broad transcriptional responses observed in steady state experiments are dominated by secondary effects. We aimed to resolve the global transcriptional dynamics ensuing after rapid deprivation of a core regulator, using MYB as an example. We performed a SLAM-seq time course experiment where genome-wide rates of nascent mRNA synthesis were serially measured following MYB degradation. First, we assessed the kinetics of global transcriptional activity by normalizing nascent mRNA counts to external spike-in controls. The first signs of global transcriptional collapse were not observed until 24 hours after MYB degradation (Fig. 4A). However, the number of affected genes increased progressively throughout the time course, and the transcriptional response continued to evolve, including altered transcription of additional core members (Fig. 4B-D). These observations are consistent with cascading secondary transcriptional changes indirectly triggered by MYB degradation. To gain additional insights into the mechanisms of secondary transcriptional collapse, we examined the total cell proteome at 1 and 2 hours post-MYB degradation. As expected, we observed profound depletion of MYB but virtually no other changes at 1 hour (Fig. 4E). In contrast, by 2 hours we saw changes in the abundance of 13 proteins, including a ~2-fold reduction of the MYC protein levels, evidently stemming from the decreased MYC transcription at the 1-hour time point (Figs. 4E, S7). This observation indicates that by 2 hours the observed transcriptional effects of MYB degradation already include significant secondary changes effected through MYC deprivation, underscoring the additional insights gained through rapid pre-steady state measurements. Indeed, gene set enrichment analysis revealed a significant downregulation of MYC targets beginning at 2 hours after MYB degradation and persisting throughout the time course, followed by a gradual downregulation of additional housekeeping pathways (Fig. 4D).

**Figure 4.**
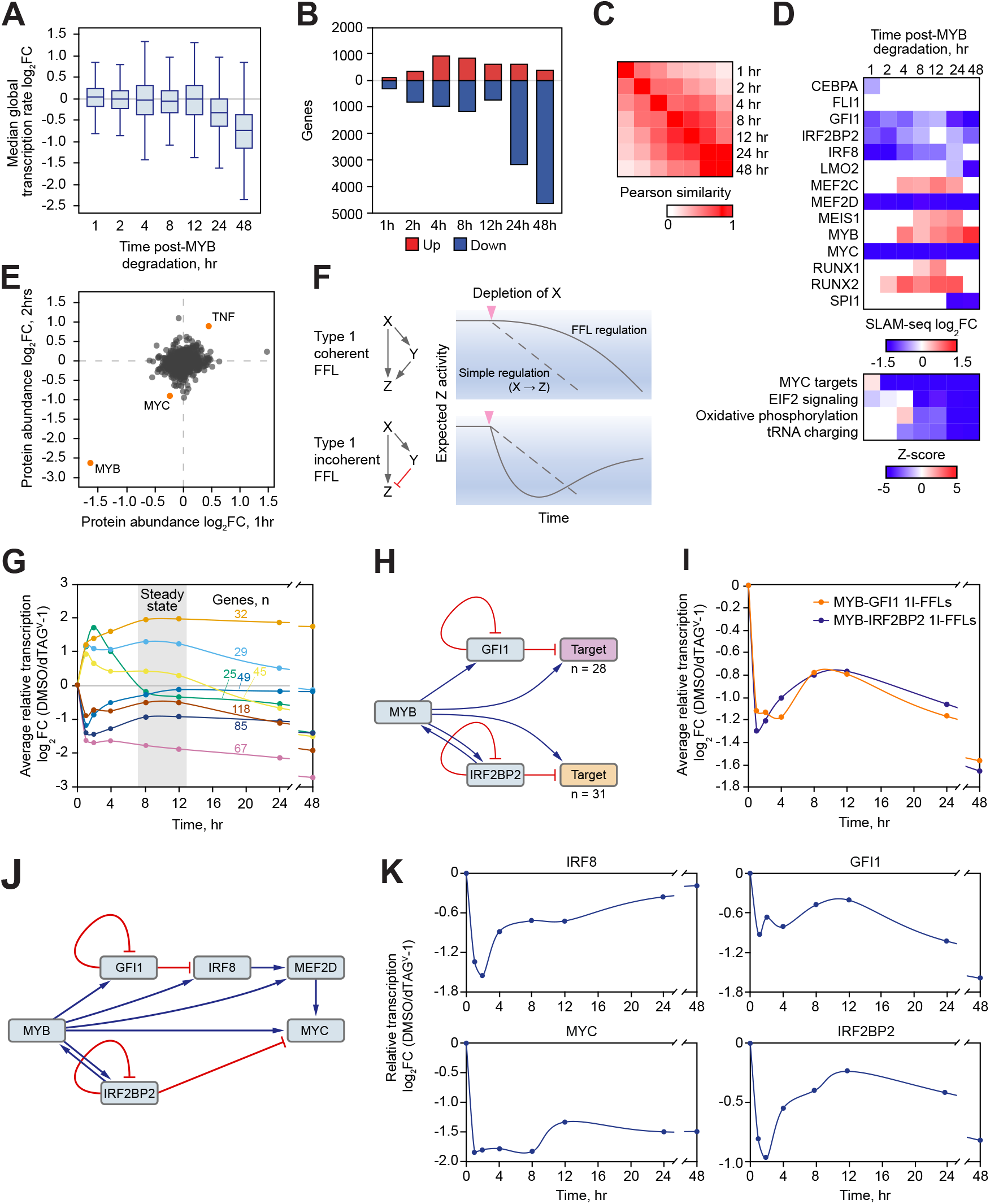
Network dynamics after MYB degradation. A, A global decrease in genome-wide transcription rates beginning at 24 hours after MYB degradation. The boxplots visualize average transcription rate changes of the top 5000 expressed genes caused by MYB degradation by dTAG^v^-1 compared to vehicle control, measured by SLAM-seq with exogenous spike-in normalization. The elements of the boxplots are as follows: center line, average log_2_ fold change in nascent transcript counts normalized to external spike-in controls; box limits, upper and lower quartiles; whiskers, 1.5x interquartile range. B, Numbers of genes demonstrating significant changes in nascent transcription at various points after MYB degradation, visualized according to the direction of response. C, Pearson similarity of the genome-scale changes in nascent transcription rates measured by SLAM-seq at various points after MYB degradation. D, Transcription rate response of the 14 core members and predicted pathway activity at various points after MYB degradation. The top heatmap visualizes log_2_ fold change in nascent transcript counts measured by SLAM-seq. Only significant (BH-adjusted *p*-value <0.05) changes are shown. The bottom heatmap visualizes predicted activation scores (z-scores) of upstream regulator pathways in Ingenuity Pathway Analysis (IPA). Data points reflect pathways with significant enrichments (BH-adjusted *p*-value <0.05, calculated by internal IPA function) and are colored according to the activity z-scores, predicting pathway inhibition (z-score <2) or activation (z-score >2). E, A scatter plot of proteome-wide changes in protein abundance detected by a quantitative proteomics experiment performed at 1 and 2 hours after MYB degradation. F, A depiction of type 1 coherent and type 1 incoherent FFLs, showing predicted dynamic behavior after the loss of an apical regulator. G, Dynamic behavior of the direct MYB target genes in a MYB degradation time course. The 450 direct MYB targets were clustered by k-means clustering by the kinetic trends of their longitudinal behavior (Pearson similarity of log_2_ fold change in transcription rates measured by SLAM-seq). Data points reflect clusteraverage log_2_ fold change in nascent transcript counts at various points after MYB degradation. H, Schematic of type 1 incoherent FFLs formed by MYB with IRF2BP2 and GFI1. I, Dynamic behavior of the genes regulated by type 1 incoherent FFLs formed by MYB with IRF2BP2 and GFI1 in a MYB degradation time course. Data points reflect average log_2_ fold change in transcription rates at various points after MYB degradation. J, Depiction of feedback and feed-forward loops in the CRC structure where MYB functions as the apical regulator. K, Dynamic behavior of the CTR genes regulated by feedback loops and type 1 incoherent FFLs originating at MYB. Data points reflect log_2_ fold change in transcription rates at various points after MYB degradation.

The feed-forward loop (FFL) is a network motif in which one regulator controls another regulator and both regulators converge on the same target gene (Fig. 4F). The regulatory effect of an FFL depends on the effect (positive vs. negative) of its edges (*36*). The canonical CRC model is based on type 1 coherent (1C)-FFLs, which have exclusive positive edges and tend to maintain a consistent level of activity resistant to transient circuit perturbations. When the apical regulator is permanently downregulated or lost, a 1C-FFL delays the response but provides no long-term buffering capacity (Fig. 4F). In contrast, FFLs with a mix of negative and positive edges, such as the type 1 incoherent (1I)-FFL, function as response accelerators, pulse generators and molecular switches (*36*). A permanently decreased activity or loss of the apical regulator elicits a bi-stable response, where a rapid primary decline in output is followed by secondary compensation and a new steady state (Fig. 4F). The presence of both positive and negative regulatory connections in the CRC structure indicates the presence of incoherent FFLs and predicts a buffering mechanism by which expression of the core regulators and their peripheral targets is restrained in steady state and re-stabilized when the system is perturbed. To test this prediction, we focused on the longitudinal transcriptional behavior of the 450 genes representing direct MYB targets. We grouped these genes into 8 clusters based on the kinetics of their transcription during the time-course, with each cluster corresponding to a distinct kinetic pattern. A significant majority of direct MYB targets demonstrated bi-phasic behavior where the initial response (positive or negative) was at least partially reversed by 8-12 hours after MYB loss, at which point the system reached a new steady state (Fig. 4G). The bistable behavior is consistent with a common presence of incoherent FFLs in the gene regulatory network. Indeed, among the 8 CTRs whose direct transcriptional targets we established, MYB forms numerous 1I-FFLs with IRF2BP2 and GFI1 (Fig. 4H). As expected, the genes regulated by these 1I-FFLs, including other core members and peripheral genes, demonstrated prominent bistable kinetics (Fig. 4I-K). We conclude that the transcriptional core itself and its extended gene-regulatory network are stabilized by incoherent FFLs. Although we could directly visualize only the FFLs formed by the 8 CTRs whose transcriptional targets we elucidated, our data suggest a common presence of incoherent FFLs in the network structure, likely formed by other transcriptional regulators not directly surveyed in this study.

### Transcriptional inhibitors act as mixed CTR agonists/antagonists

Small-molecule inhibitors of BET bromodomain proteins and CDK7 have been reported to selectively inhibit the transcriptional programs of core regulatory TFs, making them useful chemical probes for CRC studies (*37*–*39*). Two mechanisms have been proposed: direct interference with the core TF function at the target gene level (*40*, *41*), and reduced expression of the core TFs due to a disproportionate recruitment of the transcriptional apparatus to the TF-coding genes (*23*, *39*). We reasoned that benchmarking the immediate transcriptional changes induced by the inhibitors against the TF degrons would allow us to evaluate the degree and potential mechanisms of inhibitor selectivity toward the core TFs. We treated AML cells with the BET bromodomain inhibitor JQ1, the CDK7 inhibitor THZ1, and their synergistic combination (*23*) and measured the immediate transcriptional effects by SLAM-seq. JQ1 and the JQ1/THZ1 combination perturbed transcription of >2000 genes, while the direct effects of THZ1 were narrower, with 576 genes demonstrating altered transcription rates. The genome-wide effects of JQ1 and THZ1 were largely distinct from the effects of CTR degradation (Fig. 5A, S8). Nonetheless, both the core members and their peripheral direct targets were significantly enriched among inhibitor-responsive genes (Fig. 5B). Strikingly, the inhibitors displayed bimodal effects on the transcription of CTR-regulated genes, further activating a subset of genes that were repressed by CTR degradation, and vice versa, thereby acting as mixed CTR agonists/antagonists (Fig. 5C). Accordingly, although JQ1 and THZ1 perturbed transcription of most core members, expression of some CTRs was further activated, rather than inhibited (Fig. 5D). We conclude that BET bromodomain and CDK7 inhibitors display partial selectivity for the leukemia CRC by dysregulating the expression of the core members and interfering with their function at the target gene level. However, rather than phenocopying CTR loss, the inhibitors display bimodal effects, effectively acting as mixed CTR agonists/antagonists.

**Figure 5.**
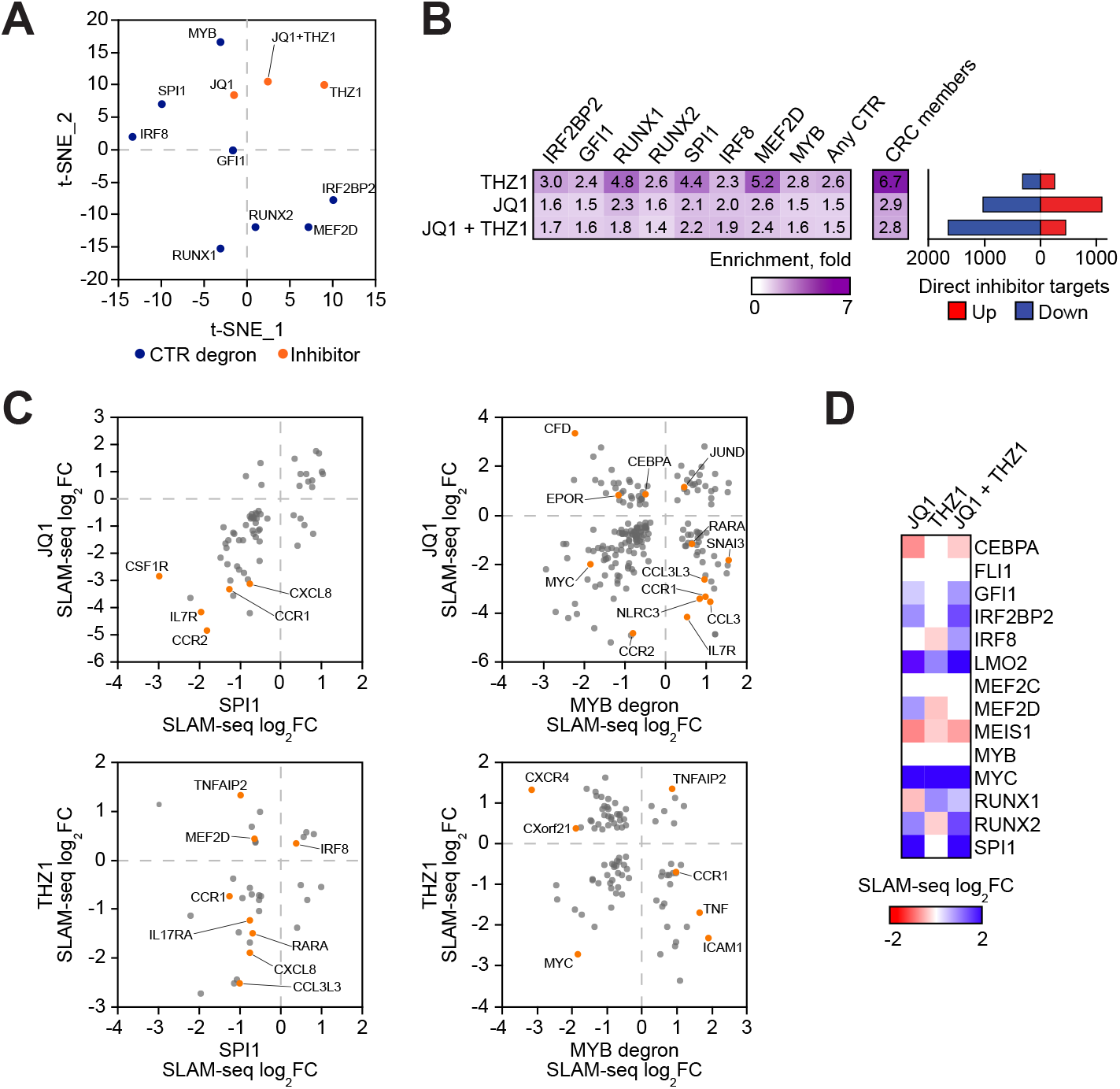
Direct transcriptional responses to BET bromodomain and CDK7 inhibition. A, A t-SNE plot of changes in the transcription rates of 5000 top expressed genes after CTR degradation and inhibitor treatment. The cells were treated for 1 hour with THZ1 (10 nM) and/or JQ1 (1 μM), versus DMSO, and the transcriptional responses were measured by SLAM-seq. B, Enrichment of the CTRs and their direct transcriptional programs among the genes whose transcription was directly affected by JQ1, THZ1 or their combination. All depicted enrichments were statistically significant (chi-square *p*-value < 0.05). C, Nascent transcriptomics responses of overlapping direct targets to CTR degradation vs. inhibitor treatment, demonstrating mixed agonist/antagonist effects of transcriptional inhibitors on the CTR-regulated genes. Data points reflect log_2_ fold change in gene transcription rates immediately after CTR degradation or 1 hour inhibitor treatment. D, Transcription rate response of the 14 core members to transcriptional inhibitors. The heatmap visualizes log_2_ fold change in nascent transcript counts after a 1 hour inhibitor treatment, measured by SLAM-seq. Only significant changes (BH-adjusted *p*-value <0.05) are shown.

### Convergence of the AML CRC on inflammation and immunity

While the extended CRC targets include genes participating in various cellular processes, we observed a strong convergence on the inflammatory and cytokine signaling pathways, likely related to the monocytic origin of the AML cells (Fig. 6A, S9). Specifically, CTRs converged on genes encoding cytokines and cytokine receptors, as well as multiple genes that modulate the activity of the immune regulator NF ºB (Fig. 6B,C). For protein-level confirmation, we assessed the levels of cytokines in the media after a 12-hour CTR depletion and performed a quantitative assessment of the proteome at the same time point after MYB degradation. In both experiments the observed protein level changes largely correlated with the nascent transcriptomics data (Fig. 6D, S9). Although at the level of individual genes the CTRs displayed pleiotropic effects (Fig. 6C), pathway analysis classified their overall regulatory vectors as either pro- or anti-inflammatory (Fig. 6A). To assess the clinical relevance of these predictions, we asked whether the expression levels of the 8 CTRs correlated with the levels of cytokines and inflammatory pathway activity in a large cohort of AML patients (*42*). The expression levels of two transcription factors, MYB and SPI1, were strongly predictive of the expression and activity of cytokines that these TFs directly regulate (Fig. 6E,F). Inflammatory pathways were enriched in comparative analyses of patients with low vs. high MYB and SPI1 expression (Fig. 6G, S10). Furthermore, we observed a strong correlation between inflammatory pathway activity changes induced by the degradation of MYB and SPI1 and the projected differences in inflammatory signaling between low- and high-expressors of these TFs among the AML patients (Fig. 6G). Notably, the pathway z-scores calculated at steady state after MYB degradation (8 hours) were more predictive of pathway activity in patients, compared to the earliest time point (1 hour) after MYB degradation (Fig. 6G). This observation further highlights the relevance of secondary events in shaping the ultimate steady state response to a change in TF level. Additional datasets, including RNA-seq of normal hematopoietic progenitors and CRISPR/Cas9 knockouts of MYB and SPI1, demonstrated a similar dependence of inflammatory signaling on the levels of MYB and SPI1 (Fig. S10). Levels of the other CTRs were less predictive of inflammatory activity in AML patients, perhaps because their overall impacts were lower and overridden by the apical regulators MYB and SPI1 (not shown).

**Figure 6.**
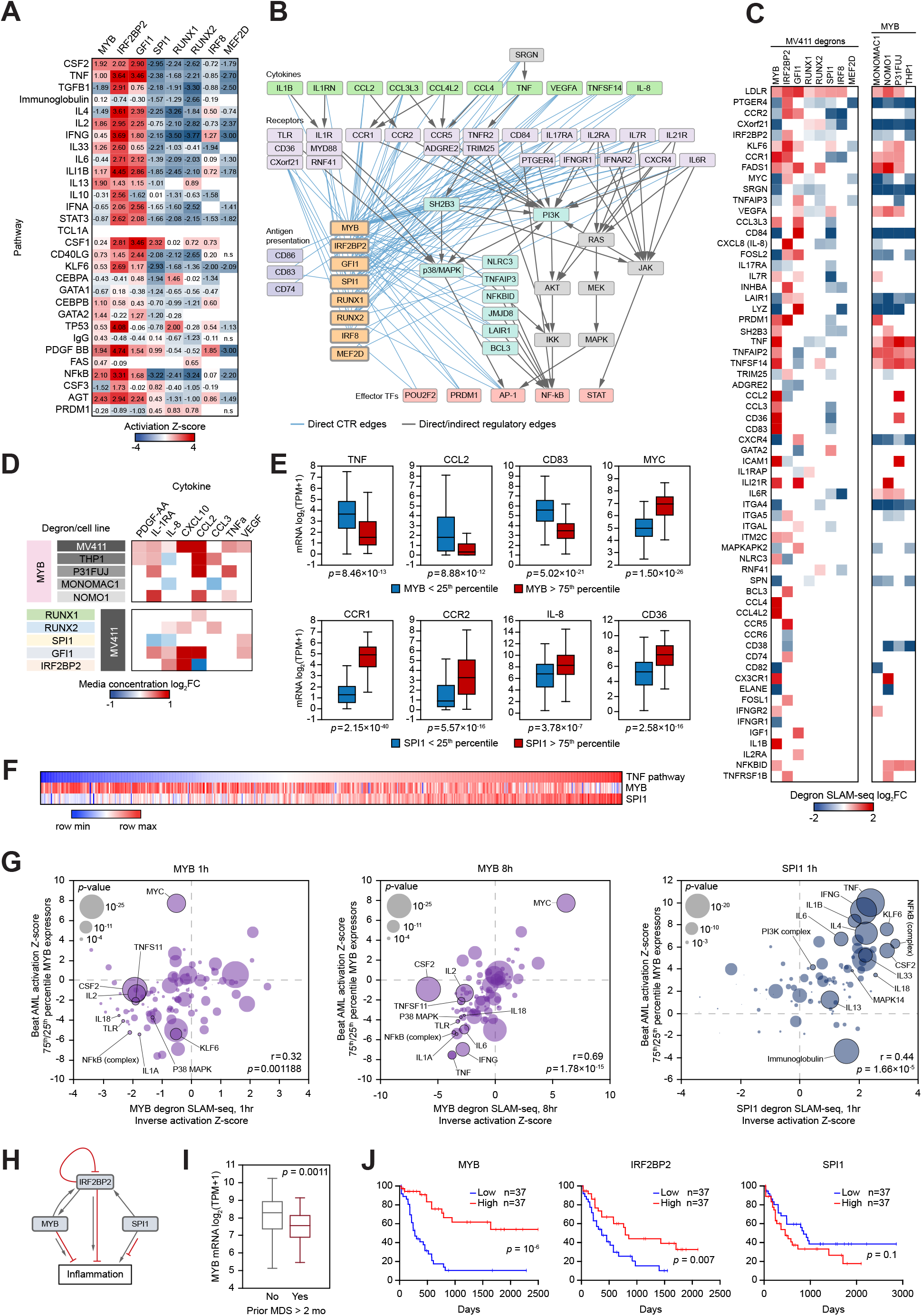
AML CRC in control of inflammation. A, A heatmap of predicted activation scores (z-scores) of top enriched upstream regulator pathways among the direct targets of the 8 CTRs, ordered by the average enrichment *p*-values. Data points reflect pathways with significant enrichments (BH-adjusted *p*-value <0.05, calculated by internal IPA function) and are colored according to the activity z-scores, predicting pathway inhibition (z-score <2) or activation (z-score >2). Empty cells designate significant pathway enrichments for which no z-score is not calculated by IPA. N.s., no significant pathway enrichment. B, A graphic depiction of the direct inflammatory pathway regulation by the AML CTRs. Each blue line represents a direct edge originating at a CTR. The regulatory relationships between the CTRs are not shown. C, A heatmap demonstrating responses of immunity/inflammation-relevant target genes to CTR degradation. Only significant responses are shown (BH-adjusted *p*-value <0.05). D, Changes in the media cytokine concentrations 12 hrs after CTR degradation. Only statistically significant changes (dTAG^v^-1 vs DMSO, 2-tailed t-test *p*-value <0.05, 4 replicates) are shown. E, Expression of MYB and SPI1 is predictive of their target gene expression levels in AML patients from the Beat AML dataset. The blue population represents patient samples with expression of MYB (top) or SPI1 (bottom) in the lowest quartile, and the red population represent patient samples with expression of MYB (top) or SPI1 (bottom) in the highest quartile. *P*-values were calculated by a Wilcoxon rank sum test. The elements of the boxplots are as follows: center line, median; box limits, upper and lower quartiles; whiskers, 1.5x interquartile range. F, Expression of MYB and SPI1 correlates with the TNF pathway activity in Beat AML patients (*n*=510), calculated as sample-average gene expression z-scores of the 46 genes in the IPA TNF pathway. G, Direct programs of MYB and SPI1 predict inflammatory pathway activity in AML patients. Depicted data points reflect top 100 upstream pathways enriched in Ingenuity Pathway Analysis (IPA enrichment *p*-value <0.05) among the direct targets of MYB and SPI1 detected by SLAM-seq after a 1 hr CTR degradation. Horizontal axis values represent inverse activation z-scores of these pathways (calculated by IPA) after 1 or 8 hours of CTR degradation. Vertical axis values represent pathway activation z-scores calculated by IPA on comparison of high expressors (highest quartile) versus low expressors (lowest quartile) of MYB and SPI1, respectively, in the Beat AML patient dataset. Bubble size illustrates the *p*-value of pathway enrichment in high versus low MYB or SPI1 expressors, respectively, among the Beat AML patients. The correlation of z-scores indicates that pathway inhibition after MYB or SPI1 degradation is strongly predictive of activation of the same pathway in patients with high mRNA expression of MYB or SPI1, respectively. Pearson correlation coefficients (r) are depicted along with their corresponding *p*-values. H, Schematic of the direct regulatory relationships between SPI1, MYB and IRF2BP2, and their overall influence on inflammation pathways in AML. I, MYB expression levels in AML patients with and without antecedent MDS in the Beat AML dataset. The *p*-value was calculated by a Wilcoxon rank sum test. J, Kaplan-Meyer plots of overall survival in AML patients from the TCGA study (*66*) with designated CTR expression above the 75th percentile (red line) and below the 25th percentile (blue line). Logrank *p*-values were calculated by multivariate Cox regression analysis in OncoLnc (*67*).

In prior work, we characterized IRF2BP2 as a critical repressor of NFκB-driven inflammation in AML cells (*43*). Here, we demonstrate that MYB forms a positive feedback loop with IRF2BP2. Despite partially antagonistic effects on their overlapping target genes (Fig. 2H), both proteins act as overall repressors of inflammation. In contrast, SPI1 is pro-inflammatory. Intriguingly, SPI1 directly activates IRF2BP2 expression, suggesting a fine-tuning mechanism by which the pro-inflammatory actions of SPI1 are dampened. Activation of inflammatory pathways has been associated with pre-leukemic conditions, such as clonal hematopoiesis of indeterminate potential (CHIP) and myelodysplastic syndrome (MDS), and with poor outcomes in AML (*44*–*53*). Indeed, we find that MYB expression is significantly lower in patients with history of MDS prior to the AML diagnosis (Fig. 6I). Conversely, high expression of MYB and IRF2BP2 is predictive of better outcomes in AML patients, while the pro-inflammatory SPI1 demonstrates an opposite trend (Fig. 6J). We conclude that the AML CRC significantly converges on inflammatory signaling, where MYB and SPI1 antagonize each other as major direct regulators of cytokine production and cell-intrinsic inflammation.

## Discussion

Dynamic rewiring of gene regulatory networks underpins normal development, malignant transformation, and tumor evolution (*6*, *11*, *17*, *19*, *39*, *54*–*59*). Orchestration of precise gene expression programs in a developing tissue while responding to environmental cues requires a gene regulatory network to balance robustness with plasticity (*60*). Our findings provide a new perspective on transcription network organization and reveal how this delicate balance may be achieved. Despite dense co-binding to the regulatory elements in the vicinity of most expressed genes, CTRs exhibit narrow and largely distinct functional transcriptional programs. Rather than forming a densely interconnected transcriptional core, master regulators are organized into a sparsely interconnected circuitry with limited self-regulatory behavior. The CRC displays a hierarchical structure with a mostly unidirectional flow of information. Transcriptional repressors and co-repressors are integral elements of this structure, forming type 1 incoherent FFLs with transcriptional activators. We propose that this topological framework integrates the opposing requirements for robustness and plasticity. Indeed, the limited functional interconnectedness and on-target cooperativity allow for regulators to be acquired or lost without perturbing the entire circuitry, while the incoherent FFLs serve as molecular switches, rapidly re-equilibrating the system to a new steady state. The hierarchical structure provides fine-tuning input to the downstream nodes without perturbing the apical master regulators. It remains to be elucidated whether these fundamental principles are conserved in other tissues and species. Nonetheless, our data indicate that dense interconnectivity is not necessary for network stability, as commonly accepted.

Although direct pharmacological manipulation of TFs remains challenging (*61*), BET bromodomain and CDK7 inhibitors have been shown to display partial selectivity toward core TFs by interfering with their expression and function (*38*, *39*, *62*). By comparing the nascent transcriptomics signatures of JQ1 and THZ1 to those of the TF degrons we found unexpectedly that these inhibitors act as mixed TF agonists/antagonists. Our findings underscore the need for further inquiries into the functional interplay between TFs and cofactors and the mechanistic aspects of cofactor inhibition in the native chromatin context.

We demonstrate that a change in the abundance of a master transcriptional regulator triggers a cascade of secondary events rendering the steady state response poorly representative of the CTR’s direct transcriptional program. On the other hand, the steady state may be a better reflection of a CTR’s ultimate physiologic significance in the network context. We illustrate this concept by an integrative analysis of the immediate and delayed responses to CTR depletion, which revealed MYB and SPI1 as major antagonistic regulators of inflammatory signaling in AML cells. We demonstrate that the expression levels of MYB and SPI1 are broadly predictive of cytokine expression and inflammatory pathway activity in AML patients. Higher expression of the oncogenic factor MYB paradoxically confers a better prognosis, possibly due to its anti-inflammatory properties and despite MYB serving as a major driver of MYC expression. This observation illustrates how complex contextual and dose-response relationships shape the balance between oncogenic and tumor suppressor properties of master lineage TFs (*63*–*65*).

In summary, through systematic interrogation of the direct gene-regulatory functions of eight master transcriptional regulators we uncovered a revised topological model of the human core transcriptional regulatory circuitry. The model is predictive of dynamic behaviors and population-level relationships between transcriptional regulators and their direct targets. The revised model more accurately captures the functional requirements for robust yet dynamic regulation of lineage gene expression. Our work provides a blueprint for functional dissection of gene regulatory networks in development and disease.

## Acknowledgements

M. Pimkin was supported by a Damon Runyon-Sohn Pediatric Cancer Fellowship and a Young Investigator Award from the Alex’s Lemonade Stand Foundation. This work was supported, in whole or in part, by research grants to M. Pimkin from the Curing Kids Cancer Foundation, When Everyone Survives Foundation, Pedals for Pediatrics, Children’s Cancer Research Fund, Children’s Leukemia Research Association, Hyundai Hope on Wheels, Kate Amato Foundation, Adenoid Cystic Carcinoma Research Foundation and Boston Children’s Hospital Office of Faculty Development. K Stegmaier was supported by NIH 5R35 CA210030 and NIH P50 CA206963. B. Nabet was supported by NCI K22 CA258805. J. M. Ellegast was supported by the Swiss National Science Foundation, the Lady Tata Memorial Trust, and the Pediatric Cancer Research Foundation, and is a Fellow of The Helen Gurley Brown Presidential Initiative (The Pussycat Foundation). N.V. Dharia was supported by the Julia’s Legacy of Hope St. Baldrick’s Foundation Fellowship. We thank Dr. R.G. Rowe for critical reading of this manuscript.

## Author Contributions

Conceptualization: SHO, MP

Computational methodology development and data analysis: JK, MP, NVD, KE, MWP, JW, AK, VRP, KS

Experiments: TH, MP, YH, JXF, JE, JME, AB, BN, FDB, RB, SM

Visualization: FDB, MWP, TH, JK, MP

Funding acquisition: SHO, MP

Supervision: SHO, MP, KS

Writing and review of the manuscript: all authors

## Supplementary Figure Legends

**Figure S1. Definition of a canonical CRC in human AML (1).**

A, Study schematic.

B, Enriched transcriptional dependencies in AML identified in our prior work (*31*). The heatmap depicts transcription factors identified as selective AML dependencies in the Broad Cancer Dependency Map project. Data points reflects probability of dependency > 0.5. Forty TF genes with dependency probability >0.5 in 3 or more AML cell lines are shown.

C, Validation of selective AML dependencies identified in the genome-wide screen. MV411 cells stably expressing Cas9 were transduced with GFP-expressing lentiviral vectors encoding gRNAs targeting the indicated genes (2 gRNAs per vector targeting the same gene). Fraction of GFP-positive cells was followed over time and normalized to the fraction of GFP-positive cells on day 2 after viral transduction.

**Figure S2. Definition of a canonical CRC in human AML (2).**

A, Overlap between recurrent AML superenhancers, human TFs and enriched AML dependencies to define the AML CRC.

B, List of CTRs identified by intersecting selective AML dependencies and recurrent AML superenhancers. The table shows gene characteristics, criteria for exclusion/inclusion and data availability. CTRs interrogated by degron models are highlighted in blue.

**Figure S3. Cooperative binding of CTRs to regulatory elements.**

A, ChIP-seq binding tracks of the CTRs (red tracks) and additional TFs and chromatin regulators (blue tracks) at the MYC superenhancer. The 2D plot represents a H3K27ac HiChIP connectivity map from our prior work (*31*) with the black box marking the location of the ChIP-seq tracks.

B, Analysis of a binary co-binding matrix created by mapping the binding peaks of the 14 validated CTRs to nucleosome-free regions detected by ATAC-seq. The plot depicts number of ATAC-seq peaks with *n* CTRs bound illustrating a progressive decline in the prevalence of binding sites with increasing number of cobinding CTRs.

C, Enrichment of co-binding sites with *n* CTRs bound compared to a model that assumes random distribution of TF binding across the nucleosome-free regions. Sites with 2-6 co-binding CTRs are negatively enriched (i.e. under-represented compared to a model of random binding distribution), while enrichment of sites with >6 CTRs binding increases progressively with increasing *n*.

**Figure S4. Genomic occupancy patterns of CTRs and other chromatin regulators.**A co-binding matrix was created by overlapping the nucleosome-free regions (ATAC-seq peaks) with the CTR binding peaks detected in ChIP-seq assays. The matrix was *k*-means clustered (*k*=100) on Pearson correlation of binary TF binding. Heatmap visualizes enrichment of CTR binding and other features by cluster. Note that the white/yellow color represents average binding (i.e. neither positive or negative enrichment) rather than no binding.

**Figure S5. Interrogation of the AML CRC by CRISPR/Cas9 knockouts.**

A, Validation of gene knockouts by Western blotting. MV411 cells were electroporated with pre-assembled Cas9/sgRNA complexes targeting the indicated genes and protein depletion was verified by Western blot 72 hours post electroporation.

B, Synthetic lethality of TF paralogs. MV411 cells were electroporated with *in vitro* assembled Cas9/sgRNA complexes targeting one or both TF paralogs as shown, and cell viability was measured relative to an AAVS1 (“safe harbor”) control by quantification of ATP pools using a luciferin-based assay.

C, Density plots depicting changes in the genome-wide H3K27ac levels after designated TF knockouts, measured by quantitative ChIP-seq using an external spike-in control. Each row visualizes spike-in normalized ChIP-seq signal around a single H3K27ac peak. Line plots visualize genome-average distribution of the H3K27ac signal.

**Figure S6. Engineering and validation of CTR degron models.**Five degrons, shown here, were engineered and validated in this work, and 2 additional degrons (IRF8 and MEF2D) were engineered and validated in our prior study (*31*).

A, Western blots demonstrating endogenous tagging of CTR loci by CRISPR-HDR and subsequent targeted degradation of the fusion protein.

B, Histograms illustrate time course experiments where CTR degradation is followed by FACS measurement of the fusion protein fluorescence.

C, Bar plots illustrating relative cell viability measured at various points after CTR degradation by quantification of ATP pools using a luciferin-based assay.

**Figure S7. Engineering and validation of additional MYB degron models, and quantitative proteomics after MYB degradation.**

A, Schematic demonstrating the tagging strategy.

B, Western blots demonstrating endogenous tagging of CTR loci by CRISPR-HDR and subsequent targeted degradation of the fusion protein.

C, Volcano plots of quantitative whole-cell proteomics at 1 hour and 2 hours after MYB degradation in the MV411 MYB-dTAG model.

D, MYC protein levels, measured by whole-cell quantitative proteomics at 1 hour and 2 hours after MYB degradation in the MV411 MYB-dTAG model. The experiment was performed in 4 replicates. Data are normalized to the DMSO control at respective time point. *P*-values were calculated by a 2-sided t-test. FDR values were calculated using Benjamini–Hochberg procedure.

**Figure S8. Comparative analyses of SLAM-seq data.**

A, Pairwise overlap of target genes, defined as genes with significant nascent transcriptional response to protein degradation or inhibition (FDR of <0.05). Data points illustrate fraction of columns overlapping with rows.

B, Pearson similarity matrix of SLAM-seq data in the space of 3475 genes directly regulated by at least one degron or inhibitor.

**Figure S9. Convergence of the AML CRC on inflammation and immunity.**

A, Gene set enrichment analysis of the 1076 genes - direct targets of at least one CTR. Analysis was performed using the Enrichr database with built-in statistical functions (*68*).

B, Correlation between changes in nascent transcription measured by SLAM-seq 8 hours after MYB degradation and protein abundance measured by a quantitative whole-proteome assay 12 hours after MYB degradation.

**Figure S10. Antagonistic functions of SPI1 and MYB in inflammation control.**

A, Pathway enrichment and activation z-scores derived from a comparative analysis of high versus low MYB and SPI1 expressors in the Beat AML dataset (*42*). The enrichment z-scores and p-values were calculated in the Ingenuity Pathway Analysis package (IPA; https://www.qiagenbioinformatics.com/products/ingenuity-pathway-analysis) using the standard “Upstream Regulator” function.

B, Transcriptome-wide changes in mRNA expression measured by RNA-seq 72 hours after a CRISPR/Cas9 knockout of MYB versus SPI1 are plotted. The TNF pathway genes (derived from IPA) are highlighted, demonstrating antagonistic regulation by MYB and SPI1.

C, Correlation between TNF expression and MYB/SPI1 expression in normal myeloid progenitors. RNA-seq data were downloaded from Ref. (*69*). Correlation (*r*) was calculated by Pearson test.

## METHODS

### METHOD DETAILS

#### Cell lines

AML cell lines were cultured in the RPMI-1640 media containing 10% fetal bovine serum and regularly tested to be free of *Mycoplasma spp.*

#### CRISPR/Cas9 gene knockouts by RNP electroporation

Synthetic modified sgRNA constructs were purchased from Synthego (Redwood City, CA). Ribonucleoprotein (RNP) assembly was performed by mixing 2-3 sgRNAs (a total of 120 pmol) with 8.5 μg recombinant Cas9 (Invitrogen A36499). The resulting RNP mix was electroporated into 0.3×10^6^ MV411 cells using a Lonza 4D Nucleofector, program DJ-100, in 20 μl Nucleocuvette strips (Lonza V4XC-2032). Cells were incubated in media for 72 hours post-electroporation before subsequent analyses. Knockout efficiency was confirmed by Western blotting and PCR amplification followed by indel analysis. A guide RNA targeting the AAVS1 “safe harbor” locus was used as a negative control. Cell viability was measured at indicated times post-electroporation using the CellTiter-Glo luminescent cell viability assay (Promega G7570).

#### CRISPR/Cas9 gene knockouts by lentiviral transduction

We used lentiviral delivery of gRNAs for initial validation of 11 arbitrarily chosen AML dependencies (Figure S7E). Lentiviral vectors encoding gRNAs (bicistronic vectors with 2 gRNAs per vector: pLV[2gRNA]-EGFP:T2A:Puro-U6>[hgRNA#1]-U6>[gRNA#2]) were purchased from VectorBuilder (Cyagen Biosciences, Santa Clara, CA) and packaged into lentivirus. Cas9-expressing AML cell lines were transduced by spinfection and the guide dropout was measured every 2 days for a total of 22 days as the percentage of GFP+ cells and normalized to the percentage of GFP+ cells 48 hours transduction.

#### Chromatin immunoprecipitation sequencing (ChIP-seq)

ChIP-seq for hematopoietic TFs was performed using 100×10^6^ exponentially growing MV411 cells per experiment. For histone ChIP-seq, 2.5×10^6^ cells were used. Cells were fixed with 1% formaldehyde for 10 minutes at room temperature, quenched with 125 mM glycine for 5 minutes, and washed 3 times with PBS. Nuclei were isolated using Nuclei EZ Isolation buffer (Sigma NUC-101) and resuspended in 10 mM Tris-HCl, pH 8.0, 1 mM EDTA, 0.1% SDS with 1x HALT protease inhibitor (ThermoFisher 78430). Chromatin was fragmented by sonication on an E220 Covaris focused sonication machine using 1 ml glass AFA tubes (Covaris 520135) and the following parameters: 140 mV, 5% duty factor, 200 cycles/burst for 14 minutes. The rest of the ChIP-seq protocol was performed as previously described (*70*) and in accordance with the Encode guidelines (*71*). ChIPseq libraries were prepared using Swift S2 Acel reagents (Swift 21096) on a Beckman Coulter Biomek i7 liquid handling platform from approximately 1 ng of DNA according to manufacturer’s protocol and using 14 cycles of PCR amplification. Sequencing libraries were quantified by Qubit fluorometer and Agilent TapeStation 2200. Library pooling and indexing was evaluated by shallow sequencing on Illumina MiSeq. Subsequently, libraries were sequenced on Illumina NextSeq 500 or NovaSeq 6000 by the Molecular Biology Core facilities at the Dana-Farber Cancer Institute. Refer to Suppl. Data for a list of native antibodies used. ChIP-seq of GFI1, IRF2BP2 and MEF2D was performed using an anti-FLAG antibody (Sigma F1804) in the MV411 strains harboring the dTAG cassette knockins which included a 3xFLAG tag (Fig. 2A) after native antibody pulldowns failed to produce data of satisfactory quality.

For quantitative ChIP-seq analysis of H3K27 acetylation and TF binding we used *Drosophila* chromatin/antibody spike-in control as previously described (*72*). For H3K27ac ChIP-seq, 4 μg of anti-H3K27ac antibody, 2 μg of spikein antibody and 20 ng of spike-in chromatin (Active Motif 61686 and 53083, respectively) were added to chromatin prepared from 2.5×10^6^ MV411 cells 72 hours after RNP-mediated TF knockout. For TF ChIP-seq, 10 μg of anti-TF antibody, 5 μg of spike-in antibody and 50 ng of spike-in chromatin were added to chromatin prepared from 100×10^6^ MV411 cells. The rest of the ChIP-seq experiment was performed in the standard fashion. After ChIP-seq reads were mapped to the *Drosophila* genome and the hg38 human genome in parallel, human tag counts were normalized to *Drosophila* tag counts.

#### RNA-seq

For RNA-seq experiments the total cellular RNA was extracted using the QuickRNA kit (Zymo Research R1054). Purified total RNA was mixed with the ERCC ExFold RNA Spike-in mix (Invitrogen 4456740). RNA sequencing libraries were prepared on a Beckman Coulter Biomek i7 liquid handling platform using Roche Kapa mRNA HyperPrep strand specific sample preparation kits (Roche 08098123702) from 200 ng of purified total RNA according to the manufacturer’s protocol. Library quantification and Illumina sequencing were performed as described in the ChIP-seq section above.

#### Western blotting

Whole-cell lysates were prepared in RIPA buffer (Boston Bio-Products BP-115-500) with protease inhibitor cocktail (ThermoFisher 23225). Lysates were boiled in Laemmli buffer (BioRad 1610737), separated by SDS-PAGE, and transferred and blocked using standard methodology. HRP-conjugated anti-mouse and anti-rabbit IgG secondary antibodies were used for imaging (BioRad 1706515 and 1706515) with an enhanced chemiluminescence substrate (PerkinElmer NEL104001EA) according to manufacturers’ instructions.

#### Targeted TF degradation

MV411 cells were modified by CRISPR-HDR to express C-terminal FKBP12 fusions of MYB, RUNX1, RUNX2, GFI1, IRF2BP2 and SPI1, respectively. Degron models of IRF8 and MEF2D had been engineered in our prior work (*31*). For each knock-in, donor DNA constructs were chemically synthesized and cloned into the pUC19 (IRF2BP2) or pAAV-MCS2 (MYB, RUNX1, RUNX2, GFI1 and SPI1) plasmids obtained from Addgene (Watertown, MA). The donors included 400-800 bp homology arms flanking the inserted DNA sequence encoding the FKBP12 degradation tag as well as mScarlet, FLAGx3 and HA tags. MV411 cells were electroporated with Cas9/sgRNA complexes targeting the HDR insertion site (with sgRNA protospacer sequence spanning the insertion site). Electroporation was performed using Lonza SF Cell Line 4D Nucleofector (V4XC-2032). RNP complexes were formed by mixing 8.5 μg of TrueCut Cas9 Protein v2 (Invitrogen A36499) and 120 pmol sgRNA. 0.3×10^6^ cells were washed with PBS and resuspended in 20 μL of SF Cell Line solution (Lonza). The cells were combined with the RNP mix and electroporated using program DJ-100. For the IRF2BP2 knockin 0.6 μg of pDNA donor was added to the RNP mix immediately prior to electroporation. A two-step process was used where the two IRF2BP2 alleles were modified sequentially using a GFP and an mScarlet donor cassette, respectively. All other knockins were carried out in one step where delivery of a single-stranded DNA HDR-donor-template was achieved by using crude preparations of recombinant adeno-associated virus (rAAV) as described (*73*). rAAV packaging was performed at the Boston Children’s Viral Core from the HDR donor constructs cloned into the pAAV-MCS2 vector. Ten μL of crude rAAV supernatant was added to the cells immediately after electroporation. After a 5-7 day incubation period the cells were sorted for mScarlet fluorescence. Single clones were then obtained by single-cell dilution microwell plating and screened for bi-allelic donor insertion by PCR. Clones were validated by Western blotting and Sanger sequencing. TF degradation was induced by adding 500 nM of dTAG^v^-1 as previously described (*34*) and followed by FACS measurement of mScarlet fluorescence and Western blotting. Time to near-complete protein degradation was empirically determined for each degron model to be as follows: IRF2BP2, 3 hours; IRF8, MEF2D and GFI1, 2 hours; MYB, RUNX1, RUNX2 and SPI1, 1 hour (Fig. S6).

#### SLAM-seq

Thiol (SH)-linked alkylation for the metabolic sequencing of RNA (SLAM-seq) was performed as described (*35*). Briefly, a total of 2.5×10^6^ MV411 cells per replicate were incubated with 500 nM dTAG^V^-1, or DMSO, for 1– 48 hours as indicated. For inhibitor experiments, MV411 cells were treated with THZ1 (10 nM) and/or JQ1 (1 μM), or DMSO, for 1 hour. All experiments were done in at least 4 replicates. S^4^U labeling was performed by adding S^4^U to a final concentration of 100 μM for an additional hour. Cells were flash-frozen and total RNA was extracted using Quick-RNA MiniPrep (Zymo Research) according to the manufacturer’s instructions except including 0.1 mM DTT to all buffers. ERCC ExFold RNA Spike-in mix (Invitrogen) was added to purified RNA. Thiol modification was performed by 10 mM iodoacetamide treatment followed by quenching with 20 mM DTT. RNA was purified by ethanol precipitation and mRNA-seq was performed as described above.

#### Cytokine measurements in culture media

Measurements of cytokine levels in culture media was done with a Luminex MagPixTM system and raw data were processes using MilliplexTM Analyst software according to manufacturer’s recommendations. The cytokine concentrations were derived from the best-fit standard curve determined by regression analysis for each cytokine. The following cytokines were measured: EGF, FGF-2, Eotaxin, TGF-α, G-CSF, FIt-3L, GM-CSF, Fractalkine, IFNα2, IFNγ, GRO, IL-10, MCP-3, IL-12p40, MDC, IL-12p70, PDGF-AA, IL-13, PDGF-AB/BB, IL-15, sCD40L, IL-17A, IL-1RA, IL-1α, IL-9, IL-1β, IL-2, IL-3, IL-4, IL-5, IL-6, IL-7, IL-8, IP-10, MCP-1, MIP-1α, MIP-1β, RANTES, TNFα, TNFβ, and VEGF. The assay was performed in four biological replicates and the difference between DMSO and dTAG^v^-1 treated samples was evaluated by a 2-sided t-test.

#### Quantitative whole-cell proteomics

Samples were prepared essentially as previously described (*74*, *75*). Following lysis, protein precipitation, reduction/alkylation and digestion, peptides were quantified by BCA assay and 150 μg of peptide per sample were labeled with TMTPro reagents (ThermoFisher) for 2 hrs at room temperature. Labeling reactions were quenched with 0.5% hydroxylamine and acidified with TFA. Acidified peptides were combined and desalted by Sep-Pak (Waters). TMT labeled peptides were solubilized in 5% ACN/10 mM ammonium bicarbonate, pH 8.0 and 300 μg of TMT labeled peptides was separated by an Agilent 300 Extend C18 column (3.5 μm particles, 4.6 mm ID and 250 mm in length). An Agilent 1260 binary pump coupled with a photodiode array (PDA) detector (ThermoFisher) was used to separate the peptides. A 45-min linear gradient from 10% to 40% acetonitrile in 10 mM ammonium bicarbonate pH 8.0 (flow rate of 0.6 mL/min) separated the peptide mixtures into a total of 96 fractions (36 seconds). A total of 96 Fractions were consolidated into 24 samples, acidified with 20 μL of 10% formic acid and vacuum dried to completion. Each sample was desalted via Stage Tips and re-dissolved in 5% formic acid/ 5% acetonitrile for LC-MS3 analysis. Proteome data were collected on an Orbitrap Eclipse mass spectrometer (ThermoFisher) coupled to a Proxeon EASY-nLC 1200 LC pump (ThermoFisher). Fractionated peptides were separated using a 180 min gradient at 500 nL/min on a 35 cm column (i.d. 100 μm, Accucore, 2.6 μm, 150 Å) packed in-house. MS1 data were collected in the Orbitrap (120,000 resolution; maximum injection time 50 ms; AGC 10 × 105). Charge states between 2 and 5 were required for MS2 analysis, and a 180 s dynamic exclusion window was used. Top 10 MS2 scans were performed in the ion trap with CID fragmentation (isolation window 0.5 Da; Turbo; NCE 35%; maximum injection time 35 ms; AGC 1.5 × 104). An on-line real-time search algorithm (Orbiter) was used to trigger MS3 scans for quantification (*76*). MS3 scans were collected in the Orbitrap using a resolution of 50,000, NCE of 55%, maximum injection time of 200 ms, and AGC of 3.0 × 105. The close out was set at two peptides per protein per fraction. Raw files were converted to mzXML, and monoisotopic peaks were re-assigned using Monocle (*77*).

Searches were performed using the Comet search algorithm against a human database downloaded from Uniprot in May 2021. We used a 50 ppm precursor ion tolerance, 1.0005 fragment ion tolerance, and 0.4 fragment bin offset for MS2 scans collected in the ion trap, and 0.02 fragment ion tolerance; 0.00 fragment bin offset for MS2 scans collected in the Orbitrap. TMTpro on lysine residues and peptide N-termini (+304.2071 Da) and carbamidomethylation of cysteine residues (+57.0215 Da) were set as static modifications, while oxidation of methionine residues (+15.9949 Da) was set as a variable modification. For phosphorylated peptide analysis, +79.9663 Da was set as a variable modification on serine, threonine, and tyrosine residues. Each run was filtered separately to 1% False Discovery Rate (FDR) on the peptide-spectrum match (PSM) level. Then proteins were filtered to the target 1% FDR level across the entire combined data set. Phosphorylation site localization was determined using the AScorePro algorithm (*78*). For reporter ion quantification, a 0.003 Da window around the theoretical m/z of each reporter ion was scanned, and the most intense m/z was used. Reporter ion intensities were adjusted to correct for isotopic impurities of the different TMTpro reagents according to manufacturer specifications. Peptides were filtered to include only those with a summed signal-to-noise (SN) ≥ 160 across all TMT channels. An extra filter of an isolation specificity (“isolation purity”) of at least 0.5 in the MS1 isolation window was applied for the phosphorylated peptide analysis. For each protein, the filtered peptide TMTpro SN values were summed to generate protein quantification values. The signal-to-noise (S/N) measurements of peptides assigned to each protein were summed. These values were normalized so that the sum of the signal for all proteins in each channel was equivalent thereby accounting for equal protein loading.

### EXTERNAL DATASETS

RNA-seq BAM files for the BeatAML project (*42*) were provided by Oregon Health & Science University and processed through the CCLE RNA processing pipeline (STAR/RSEM, described at https://github.com/broadinstitute/ccle_processing). Reads were normalized to transcripts per million (TPM) and filtered for protein coding genes. The expression values were transformed to log2(TPM+1).

Genetic dependency data are available for download at the Broad DepMap portal database (https://depmap.org/portal/download/). Data release 20q1 was used for this study.

ATAC-seq data in MV411 cells were downloaded from https://www.ncbi.nlm.nih.gov/sra?term=SRX5608489.

Details of additional external ChIP-seq datasets are found in Supplementary Data 1.

### QUANTIFICATION AND STATISTICAL ANALYSIS DETAILS

#### ChIP-seq data analysis

Quality control, mapping and analysis of the ChIP-seq data was performed using the nf-core pipeline (https://github.com/nf-core/chipseq). Differential binding of the same protein under two conditions was computed using the *diffpeak* function of the MACS2 pipeline (https://github.com/macs3-project/MACS). Spike-in controlled experiments were mapped to the *Drosophila* genome and the hg38 human genome in parallel, and human tag counts were normalized to *Drosophila* tag counts as described.

#### Merging of TF ChIP-seq replicates

We developed CREME, a fast post-processing ChIP-seq replicate merging algorithm that does not need any raw sequencing data. CREME uses as input sets of BED files and bigWig tracks for each protein, and outputs a merged BED file, as well as merging quality metrics. Given a set of replicates, CREME first computes a consensus set of peaks by taking the union of all of their peaks and considering any peak ≤150 bp away from another to be in overlap. The overlapping peaks are then merged and assigned the mean of their signals and the product of their p-values. CREME then computes a similarity score as follows:

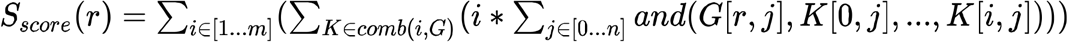

where:

*G* is an *m*n* binary matrix of *m* replicates with *n* consensus peaks and a value of 1 if replicate *m_i_*, has a peak on consensus peak *m_i_*, and 0 otherwise,

*r* is one of the replicates,

*combi(i, G)* is a list of all possible matrices made from taking *i* replicates from matrix *G* without replacement,

*and(*) is a binary operation returning 1 if all passed elements are 1 else 0.

The highest scoring sample not labelled as a poor quality replicate will be selected as the main replicate (replicate A). Poor quality replicates are user provided annotations, *e.g.* found by visual inspection of bigWig tracks, a threshold on FRiP scores or any other QC method. If the second best scoring replicate (replicate B) and the main replicate have less than 30% peak overlap CREME marks that ChIP as failed and only returns the main replicate.

Then, for each replicate, CREME searches for new peaks using a modified MACS2 peak calling algorithm with a lower discovery threshold (a KL distance of 8). We assume that the peaks are ‘events’ in a Poisson process occurring across the genome and compute the difference between the region of interest and its entire chromosome, using the KL divergence between two poisson distributions originating from these two regions. The signal for these events is extracted from the bigWig files. If the KL distance is above the threshold we consider the region as a peak. Using this approach, CREME will attempt to call peaks in replicate B using peak loci that were called in replicate A and do not overlap with replicate B. The same is then repeated in reverse, attempting to call peaks in replicate A using non-overlapping peak regions from replicate B. If, despite the additional peak calling, we still detect less than 30% overlap between replicates, replicate B is discarded. Otherwise, the newly called peaks are added to the merged peak set. CREME then repeats this procedure with additional replicates ranked in the order of their quality. The final merged BED file contains all consensus peaks of the main replicates (including its newly found peaks) and any other peaks present in at least 2 replicates. CREME source code is available at https://github.com/broadinstitute/genepy/blob/master/genepy/epigenetics/CREME.py. A more detailed description of CREME is available at https://github.com/broadinstitute/genepy/blob/master/genepy/epigenetics/CREME.md.

#### RNA-seq data analysis

RNA-seq data were processed using the CCLE workflow of STAR (v2.6.1c) + RSEM (v1.3.0) with hg38 reference genome, enriched with ERCC92 v29 reference (https://storage.googleapis.com/ccle_default_params/Homo_sapiens_assembly38_ERCC92.fasta; https://storage.googleapis.com/ccle_default_params/STAR_genome_GRCh38_noALT_noHLA_noDecoy_ERCC_v29_oh100.tar.gz;

https://storage.googleapis.com/ccle_default_params/rsem_reference_GRCh38_gencode29_ercc.tar.gz) and Gencode v29 reference gene regions (https://storage.googleapis.com/ccle_default_params/references_gtex_gencode.v29.GRCh38.ERCC.genes.collapsed_only.gtf). Scaling factors were computed from ERCC spike-ins using the ERCCdashboard R package (https://bioconductor.org/packages/release/bioc/html/erccdashboard.html). Scaling was applied only to samples where ERCC displayed a mean scaling of at least twice the size of standard error. Differential analysis of RNA-seq data was performed using DESeq2 (v1.26.0). (https://bioconductor.org/packages/release/bioc/html/DESeq2.html), using the ERCC pseudogenes to rescale the data by using *the run_estimate_size_factors control_genes* parameter.

#### SLAM-seq data analysis

A modified version of the slamdunk pipeline was used for SLAM-seq processing (available at https://github.com/jkobject/slamdunk). Differential analysis of SLAM-seq data was performed using DESeq2 on the TC converted transcripts (*tccounts*) and total read counts (*totalcounts*). First, mean *totalcounts* were used to compute scaling factors via the DESeq2 *run_estimate_size_factors geoMeans* parameter. Then, the *tccounts* and *totalcounts* were normalized to the ERCC pseudogene counts using DEseq2 *getSizeFactors* and *setSizeFactors* functions. Further details and code are found at https://github.com/jkobject/AMLproject/blob/master/notebooks/slamseq_iBet_spikeIn_maxp_paper.ipynb

#### TF co-occupancy map

To compute a TF co-occupancy map we used the genepy *simpleMergePeaks* function (https://github.com/broadinstitute/genepy/blob/master/genepy/epigenetics/chipseq.py) to merge CREME consensus peaks of the 14 CTRs. The peaks were merged if they were <150 bp apart (from edge to edge). Further details and analysis pipeline are found at https://github.com/jkobject/AMLproject/blob/master/notebooks/CoBinding%20pre-processing%20and%20analysis.ipynb and https://github.com/jkobject/AMLproject/blob/master/notebooks/CoBinding%20Enrichment%20and%20Correlation.ipynb

## References

1. K. Takahashi, S. Yamanaka, Induction of Pluripotent Stem Cells from Mouse Embryonic and Adult Fibroblast Cultures by Defined Factors. Cell. 126, 663–676 (2006).

2. T. Graf, T. Enver, Forcing cells to change lineages. Nature. 462, 587–594 (2009).

3. R. L. Davis, H. Weintraub, A. B. Lassar, Expression of a single transfected cDNA converts fibroblasts to myoblasts. Cell. 51, 987–1000 (1987).

4. S. H. Orkin, K. Hochedlinger, Chromatin Connections to Pluripotency and Cellular Reprogramming. Cell. 145, 835–850 (2011).

5. L. A. Boyer, T. I. Lee, M. F. Cole, S. E. Johnstone, S. S. Levine, J. P. Zucker, M. G. Guenther, R. M. Kumar, H. L. Murray, R. G. Jenner, D. K. Gifford, D. A. Melton, R. Jaenisch, R. A. Young, Core transcriptional regulatory circuitry in human embryonic stem cells. Cell. 122, 947–956 (2005).

6. M. A. Kerényi, S. H. Orkin, Networking erythropoiesis. The Journal of experimental medicine. 207, 2537–2541 (2010).

7. S. H. Orkin, J. Wang, J. Kim, J. Chu, S. Rao, T. W. Theunissen, X. Shen, D. N. Levasseur, The Transcriptional Network Controlling Pluripotency in ES Cells. Cold Spring Harb Sym. 73, 195–202 (2008).

8. J. Joung, S. Ma, T. Tay, K. R. Geiger-Schuller, P. C. Kirchgatterer, V. K. Verdine, B. Guo, M. A. Arias-Garcia, W. E. Allen, A. Singh, O. Kuksenko, O. O. Abudayyeh, J. S. Gootenberg, Z. Fu, R. K. Macrae, J. D. Buenrostro, A. Regev, F. Zhang, A transcription factor atlas of directed differentiation. Cell. 186, 209–229.e26 (2023).

9. T. R. Sorrells, A. D. Johnson, Making Sense of Transcription Networks. Cell. 161, 714–723 (2015).

10. E. H. Davidson, Emerging properties of animal gene regulatory networks. Nature. 468, 911–920 (2010).

11. A. C. Wilkinson, H. Nakauchi, B. Göttgens, Mammalian Transcription Factor Networks: Recent Advances in Interrogating Biological Complexity. Cell Syst. 5, 319–331 (2017).

12. S. A. Lambert, A. Jolma, L. F. Campitelli, P. K. Das, Y. Yin, M. Albu, X. Chen, J. Taipale, T. R. Hughes, M. T. Weirauch, The Human Transcription Factors. Cell. 172, 650–665 (2018).

13. W. Deng, J. Lee, H. Wang, J. Miller, A. Reik, P. D. Gregory, A. Dean, G. A. Blobel, Controlling Long-Range Genomic Interactions at a Native Locus by Targeted Tethering of a Looping Factor. Cell. 149, 1233–1244 (2012).

14. N. Liu, V. V. Hargreaves, Q. Zhu, J. V. Kurland, J. Hong, W. Kim, F. Sher, C. Macias-Trevino, J. M. Rogers, R. Kurita, Y. Nakamura, G.-C. Yuan, D. E. Bauer, J. Xu, M. L. Bulyk, S. H. Orkin, Direct Promoter Repression by BCL11A Controls the Fetal to Adult Hemoglobin Switch. Cell. 173, 430–442.e17 (2018).

15. M. R. Mansour, B. J. Abraham, L. Anders, A. Berezovskaya, A. Gutierrez, A. D. Durbin, J. Etchin, L. Lawton, S. E. Sallan, L. B. Silverman, M. L. Loh, S. P. Hunger, T. Sanda, R. A. Young, A. T. Look, Oncogene regulation. An oncogenic super-enhancer formed through somatic mutation of a noncoding intergenic element. Science. 346, 1373–1377 (2014).

16. M. G. Jaeger, G. E. Winter, Fast-acting chemical tools to delineate causality in transcriptional control. Mol Cell. 81, 1617–1630 (2021).

17. N. Novershtern, A. Subramanian, L. N. Lawton, R. H. Mak, W. N. Haining, M. E. McConkey, N. Habib, N. Yosef, C. Y. Chang, T. Shay, G. M. Frampton, A. C. B. Drake, I. Leskov, B. Nilsson, F. Preffer, D. Dombkowski, J. W. Evans, T. Liefeld, J. S. Smutko, J. Chen, N. Friedman, R. A. Young, T. R. Golub, A. Regev, B. L. Ebert, Densely interconnected transcriptional circuits control cell states in human hematopoiesis. Cell. 144, 296–309 (2011).

18. T. F. Consortium, O. J. L. Rackham, J. Firas, H. Fang, M. E. Oates, M. L. Holmes, A. S. Knaupp, H. Suzuki, C. M. Nefzger, C. O. Daub, J. W. Shin, E. Petretto, A. R. R. Forrest, Y. Hayashizaki, J. M. Polo, J. Gough, A predictive computational framework for direct reprogramming between human cell types. Nat Genet.48, 331–335 (2016).

19. S. Neph, A. B. Stergachis, A. Reynolds, R. Sandstrom, E. Borenstein, J. A. Stamatoyannopoulos, Circuitry and dynamics of human transcription factor regulatory networks. Cell. 150, 1274–1286 (2012).

20. J. Kim, J. Chu, X. Shen, J. Wang, S. H. Orkin, An Extended Transcriptional Network for Pluripotency of Embryonic Stem Cells. Cell. 132, 1049–1061 (2008).

21. C. J. Nobile, E. P. Fox, J. E. Nett, T. R. Sorrells, Q. M. Mitrovich, A. D. Hernday, B. B. Tuch, D. R. Andes, A. D. Johnson, A Recently Evolved Transcriptional Network Controls Biofilm Development in Candida albicans. Cell. 148, 126–138 (2012).

22. V. Saint-André, A. J. Federation, C. Y. Lin, B. J. Abraham, J. Reddy, T. I. Lee, J. E. Bradner, R. A. Young, Models of human core transcriptional regulatory circuitries. Genome Res. 26, 385–396 (2016).

23. A. D. Durbin, M. W. Zimmerman, N. V. Dharia, B. J. Abraham, A. B. Iniguez, N. Weichert-Leahey, S. He, J. M. Krill-Burger, D. E. Root, F. Vazquez, A. Tsherniak, W. C. Hahn, T. R. Golub, R. A. Young, A. T. Look, K. Stegmaier, Selective gene dependencies in MYCN-amplified neuroblastoma include the core transcriptional regulatory circuitry. Nat Genet. 50, 1240–1246 (2018).

24. C. J. Ott, A. J. Federation, L. S. Schwartz, S. Kasar, J. L. Klitgaard, R. Lenci, Q. Li, M. Lawlor, S. M. Fernandes, A. Souza, D. Polaski, D. Gadi, M. L. Freedman, J. R. Brown, J. E. Bradner, Enhancer Architecture and Essential Core Regulatory Circuitry of Chronic Lymphocytic Leukemia. Cancer Cell. 34, 982–995.e7 (2018).

25. Y. Chen, L. Xu, R. Y.-T. Lin, M. Muschen, H. P. Koeffler, Core transcriptional regulatory circuitries in cancer. Oncogene, 1–14 (2020).

26. M. Huang, Y. Chen, M. Yang, A. Guo, Y. Xu, L. Xu, H. P. Koeffler, dbCoRC: a database of core transcriptional regulatory circuitries modeled by H3K27ac ChIP-seq signals. Nucleic Acids Res. 46 (2017), doi:10.1093/nar/gkx796.

27. V. Boeva, C. Louis-Brennetot, A. Peltier, S. Durand, C. Pierre-Eugène, V. Raynal, H. C. Etchevers, S. Thomas, A. Lermine, E. Daudigeos-Dubus, B. Geoerger, M. F. Orth, T. G. P. Grünewald, E. Diaz, B. Ducos, D. Surdez, A. M. Carcaboso, I. Medvedeva, T. Deller, V. Combaret, E. Lapouble, G. Pierron, S. Grossetête-Lalami, S. Baulande, G. Schleiermacher, E. Barillot, H. Rohrer, O. Delattre, I. Janoueix-Lerosey, Heterogeneity of neuroblastoma cell identity defined by transcriptional circuitries. Nat Genet. 2, 16078 (2017).

28. C. Y. Lin, S. Erkek, Y. Tong, L. Yin, A. J. Federation, M. Zapatka, P. Haldipur, D. Kawauchi, T. Risch, H.-J. Warnatz, B. C. Worst, B. Ju, B. A. Orr, R. Zeid, D. R. Polaski, M. Segura-Wang, S. M. Waszak, D. T. W. Jones, M. Kool, V. Hovestadt, I. Buchhalter, L. Sieber, P. Johann, L. Chavez, S. Gröschel, M. Ryzhova, A. Korshunov, W. Chen, V. V. Chizhikov, K. J. Millen, V. Amstislavskiy, H. Lehrach, M.-L. Yaspo, R. Eils, P. Lichter, J. O. Korbel, S. M. Pfister, J. E. Bradner, P. A. Northcott, Active medulloblastoma enhancers reveal subgroup-specific cellular origins. Nature. 530, 57–62 (2016).

29. M. Muhar, A. Ebert, T. Neumann, C. Umkehrer, J. Jude, C. Wieshofer, P. Rescheneder, J. J. Lipp, V. A. Herzog, B. Reichholf, D. A. Cisneros, T. Hoffmann, M. F. Schlapansky, P. Bhat, A. von Haeseler, T. Köcher, A. C. Obenauf, J. Popow, S. L. Ameres, J. Zuber, SLAM-seq defines direct gene-regulatory functions of the BRD4-MYC axis. Science. 360, 800–805 (2018).

30. K. R. Stengel, J. D. Ellis, C. L. Spielman, M. L. Bomber, S. W. Hiebert, Definition of a small core transcriptional circuit regulated by AML1-ETO. Mol Cell. 81, 530–545.e5 (2021).

31. T. Harada, Y. Heshmati, J. Kalfon, M. W. Perez, J. X. Ferrucio, J. Ewers, B. H. Engler, A. Kossenkov, J. M. Ellegast, J. S. Yi, A. Bowker, Q. Zhu, K. Eagle, T. Liu, Y. Kai, J. M. Dempster, G. Kugener, J. Wickramasinghe, Z. T. Herbert, C. H. Li, J. V. Koren, D. M. Weinstock, V. R. Paralkar, B. Nabet, C. Y. Lin, N. V. Dharia, K. Stegmaier, S. H. Orkin, M. Pimkin, A distinct core regulatory module enforces oncogene expression in KMT2A-rearranged leukemia. Gene Dev. 36, 368–389 (2022).

32. K. Chen, Z. Hu, Z. Xia, D. Zhao, W. Li, J. K. Tyler, The Overlooked Fact: Fundamental Need for Spike-In Control for Virtually All Genome-Wide Analyses. Molecular and cellular biology. 36, 662–667 (2015).

33. J. Lovén, D. A. Orlando, A. A. Sigova, C. Y. Lin, P. B. Rahl, C. B. Burge, D. L. Levens, T. I. Lee, R. A. Young, Revisiting global gene expression analysis. Cell. 151, 476–482 (2012).

34. B. Nabet, F. M. Ferguson, B. K. A. Seong, M. Kuljanin, A. L. Leggett, M. L. Mohardt, A. Robichaud, A. S. Conway, D. L. Buckley, J. D. Mancias, J. E. Bradner, K. Stegmaier, N. S. Gray, Rapid and direct control of target protein levels with VHL-recruiting dTAG molecules. Nature communications. 11, 4687–8 (2020).

35. V. A. Herzog, B. Reichholf, T. Neumann, P. Rescheneder, P. Bhat, T. R. Burkard, W. Wlotzka, A. von Haeseler, J. Zuber, S. L. Ameres, Thiol-linked alkylation of RNA to assess expression dynamics. Nat Methods.14, 1198–1204 (2017).

36. U. Alon, Network motifs: theory and experimental approaches. Nature reviews. Genetics. 8, 450–461 (2007).

37. B. E. Gryder, L. Wu, G. M. Woldemichael, S. Pomella, T. R. Quinn, P. M. C. Park, A. Cleveland, B. Z. Stanton, Y. Song, R. Rota, O. Wiest, M. E. Yohe, J. F. Shern, J. Qi, J. Khan, Chemical genomics reveals histone deacetylases are required for core regulatory transcription. Nature communications. 10, 3004–12 (2019).

38. S. Sengupta, R. E. George, Super-Enhancer-Driven Transcriptional Dependencies in Cancer. Trends in cancer. 3, 269–281 (2017).

39. J. E. Bradner, D. Hnisz, R. A. Young, Transcriptional Addiction in Cancer. Cell. 168, 629–643 (2017).

40. J.-S. Roe, F. Mercan, K. Rivera, D. J. Pappin, C. R. Vakoc, BET Bromodomain Inhibition Suppresses the Function of Hematopoietic Transcription Factors in Acute Myeloid Leukemia. Mol Cell. 58, 1028–1039 (2015).

41. A. S. Bhagwat, J.-S. Roe, B. Y. L. Mok, A. F. Hohmann, J. Shi, C. R. Vakoc, BET Bromodomain Inhibition Releases the Mediator Complex from Select cis-Regulatory Elements. Cell Reports. 15, 519–530 (2016).

42. J. W. Tyner, C. E. Tognon, D. Bottomly, B. Wilmot, S. E. Kurtz, S. L. Savage, N. Long, A. R. Schultz, E. Traer, M. Abel, A. Agarwal, A. Blucher, U. Borate, J. Bryant, R. Burke, A. Carlos, R. Carpenter, J. Carroll, B. H. Chang, C. Coblentz, A. d’Almeida, R. Cook, A. Danilov, K.-H. T. Dao, M. Degnin, D. Devine, J. Dibb, D. K. Edwards, C. A. Eide, I. English, J. Glover, R. Henson, H. Ho, A. Jemal, K. Johnson, R. Johnson, B. Junio, A. Kaempf, J. Leonard, C. Lin, S. Q. Liu, P. Lo, M. M. Loriaux, S. Luty, T. Macey, J. MacManiman, J. Martinez, M. Mori, D. Nelson, C. Nichols, J. Peters, J. Ramsdill, A. Rofelty, R. Schuff, R. Searles, E. Segerdell, R. L. Smith, S. E. Spurgeon, T. Sweeney, A. Thapa, C. Visser, J. Wagner, K. Watanabe-Smith, K. Werth, J. Wolf, L. White, A. Yates, H. Zhang, C. R. Cogle, R. H. Collins, D. C. Connolly, M. W. Deininger, L. Drusbosky, C. S. Hourigan, C. T. Jordan, P. Kropf, T. L. Lin, M. E. Martinez, B. C. Medeiros, R. R. Pallapati, D. A. Pollyea, R. T. Swords, J. M. Watts, S. J. Weir, D. L. Wiest, R. M. Winters, S. K. McWeeney, B. J. Druker, Functional genomic landscape of acute myeloid leukaemia. Nature. 562, 526–531 (2018).

43. J. M. Ellegast, G. Alexe, A. Hamze, S. Lin, H. J. Uckelmann, P. J. Rauch, M. Pimkin, L. S. Ross, N. V. Dharia, A. L. Robichaud, A. S. Conway, D. Khalid, J. A. Perry, M. Wunderlich, L. Benajiba, Y. Pikman, B. Nabet, N. S. Gray, S. H. Orkin, K. Stegmaier, Unleashing cell-intrinsic inflammation as a strategy to kill AML blastsCell-intrinsic inflammation as a strategy to kill AML blasts. Cancer Discov (2022), doi:10.1158/2159-8290.cd-21-0956.

44. S. Avagyan, J. E. Henninger, W. P. Mannherz, M. Mistry, J. Yoon, S. Yang, M. C. Weber, J. L. Moore, L. I. Zon, Resistance to inflammation underlies enhanced fitness in clonal hematopoiesis. Science. 374, 768–772 (2021).

45. D. A. Sallman, A. List, The central role of inflammatory signaling in the pathogenesis of myelodysplastic syndromes. Blood. 133, 1039–1048 (2019).

46. S. Binder, M. Luciano, J. Horejs-Hoeck, The Cytokine Network in Acute Myeloid Leukemia (AML): A Focus on Pro-and Anti-Inflammatory Mediators. Cytokine Growth F R. 43, 8–15 (2018).

47. A. Yeaton, G. Cayanan, S. Loghavi, I. Dolgalev, E. M. Leddin, C. E. Loo, H. Torabifard, D. Nicolet, J. Wang, K. Corrigan, V. Paraskevopoulou, D. T. Starczynowski, E. Wang, O. Abdel-Wahab, A. D. Viny, R. M. Stone, J. C. Byrd, O. A. Guryanova, R. M. Kohli, G. A. Cisneros, A. Tsirigos, A.-K. Eisfeld, I. Aifantis, M. Guillamot, The Impact of Inflammation-Induced Tumor Plasticity during Myeloid Transformation. Cancer Discov. 12, 2392–2413 (2022).

48. Y. Kagoya, A. Yoshimi, K. Kataoka, M. Nakagawa, K. Kumano, S. Arai, H. Kobayashi, T. Saito, Y. Iwakura, M. Kurokawa, Positive feedback between NF-κB and TNF-α promotes leukemia-initiating cell capacity. J Clin Invest. 124, 528–542 (2014).

49. S. Stratmann, S. A. Yones, M. Garbulowski, J. Sun, A. Skaftason, M. Mayrhofer, N. Norgren, M. K. Herlin, C. Sundström, A. Eriksson, M. Höglund, J. Palle, J. Abrahamsson, K. Jahnukainen, M. C. Munthe-Kaas, B. Zeller, K. P. Tamm, L. Cavelier, J. Komorowski, L. Holmfeldt, Transcriptomic analysis reveals pro-inflammatory signatures associated with acute myeloid leukemia progression. Blood Adv. 6, 152–164 (2021).

50. M. Yamashita, E. Passegué, TNF-α Coordinates Hematopoietic Stem Cell Survival and Myeloid Regeneration. Cell Stem Cell. 25, 357–372.e7 (2019).

51. S. O. Abegunde, R. Buckstein, R. A. Wells, M. J. Rauh, An inflammatory environment containing TNFα favors Tet2-mutant clonal hematopoiesis. Exp Hematol. 59, 60–65 (2018).

52. J. J. Trowbridge, D. T. Starczynowski, Innate immune pathways and inflammation in hematopoietic aging, clonal hematopoiesis, and MDS. J Exp Med. 218, e20201544 (2021).

53. A. Lasry, B. Nadorp, M. Fornerod, D. Nicolet, H. Wu, C. J. Walker, Z. Sun, M. T. Witkowski, A. N. Tikhonova, M. Guillamot-Ruano, G. Cayanan, A. Yeaton, G. Robbins, E. A. Obeng, A. Tsirigos, R. M. Stone, J. C. Byrd, S. Pounds, W. L. Carroll, T. A. Gruber, A.-K. Eisfeld, I. Aifantis, An inflammatory state remodels the immune microenvironment and improves risk stratification in acute myeloid leukemia. Nat Cancer, 1–16 (2022).

54. K. Eagle, T. Harada, J. Kalfon, M. W. Perez, Y. Heshmati, J. Ewers, J. V. Koren, J. M. Dempster, G. Kugener, V. R. Paralkar, C. Y. Lin, N. V. Dharia, K. Stegmaier, S. H. Orkin, M. Pimkin, Transcriptional plasticity drives leukemia immune escape. Blood Cancer Discov (2022), doi:10.1158/2643-3230.bcd-21-0207.

55. G. May, S. Soneji, A. J. Tipping, J. Teles, S. J. McGowan, M. Wu, Y. Guo, C. Fugazza, J. Brown, G. Karlsson, C. Pina, V. Olariu, S. Taylor, D. G. Tenen, C. Peterson, T. Enver, Dynamic analysis of gene expression and genome-wide transcription factor binding during lineage specification of multipotent progenitors. Cell Stem Cell. 13, 754–768 (2013).

56. A. M. Tsankov, H. Gu, V. Akopian, M. J. Ziller, J. Donaghey, I. Amit, A. Gnirke, A. Meissner, Transcription factor binding dynamics during human ES cell differentiation. Nature. 518, 344–349 (2015).

57. S. A. Assi, M. R. Imperato, D. J. L. Coleman, A. Pickin, S. Potluri, A. Ptasinska, P. S. Chin, H. Blair, P. Cauchy, S. R. James, J. Zacarias-Cabeza, L. N. Gilding, A. Beggs, S. Clokie, J. C. Loke, P. Jenkin, A. Uddin, R. Delwel, S. J. Richards, M. Raghavan, M. J. Griffiths, O. Heidenreich, P. N. Cockerill, C. Bonifer, Subtype-specific regulatory network rewiring in acute myeloid leukemia. Nat Genet. 51, 1 (2018).

58. D. K. Goode, N. Obier, M. S. Vijayabaskar, M. Lie-A-Ling, A. J. Lilly, R. Hannah, M. Lichtinger, K. Batta, M. Florkowska, R. Patel, M. Challinor, K. Wallace, J. Gilmour, S. A. Assi, P. Cauchy, M. Hoogenkamp, D. R. Westhead, G. Lacaud, V. Kouskoff, B. Göttgens, C. Bonifer, Dynamic Gene Regulatory Networks Drive Hematopoietic Specification and Differentiation. Dev Cell. 36, 572–587 (2016).

59. J. I. Sive, B. Göttgens, Transcriptional network control of normal and leukaemic haematopoiesis. Exp Cell Res. 329, 255–264 (2014).

60. L. T. MacNeil, A. J. M. Walhout, Gene regulatory networks and the role of robustness and stochasticity in the control of gene expression. Genome Res. 21, 645–657 (2011).

61. M. J. Henley, A. N. Koehler, Advances in targeting ‘undruggable’ transcription factors with small molecules. Nat Rev Drug Discov. 20, 669–688 (2021).

62. S. J. Vervoort, J. R. Devlin, N. Kwiatkowski, M. Teng, N. S. Gray, R. W. Johnstone, Targeting transcription cycles in cancer. Nat Rev Cancer. 22, 5–24 (2022).

63. N. Datta, S. Chakraborty, M. Basu, M. K. Ghosh, Tumor Suppressors Having Oncogenic Functions: The Double Agents. Cells. 10, 46 (2020).

64. M. L. Clarke, R. B. Lemma, D. S. Walton, G. Volpe, B. Noyvert, O. S. Gabrielsen, J. Frampton, MYB insufficiency disrupts proteostasis in hematopoietic stem cells leading to age-related neoplasia. Blood (2023), doi:10.1182/blood.2022019138.

65. P. García, M. Clarke, A. Vegiopoulos, O. Berlanga, A. Camelo, M. Lorvellec, J. Frampton, Reduced c□Mvb activity compromises HSCs and leads to a myeloproliferation with a novel stem cell basis. Embo J. 28, 1492–1504 (2009).

66. C. G. A. R. Network, Genomic and epigenomic landscapes of adult de novo acute myeloid leukemia. New Engl J Medicine. 368, 2059–2074 (2013).

67. J. Anaya, OncoLnc: linking TCGA survival data to mRNAs, miRNAs, and lncRNAs. Peerj Comput Sci. 2, e67 (2016).

68. M. V. Kuleshov, M. R. Jones, A. D. Rouillard, N. F. Fernandez, Q. Duan, Z. Wang, S. Koplev, S. L. Jenkins, K. M. Jagodnik, A. Lachmann, M. G. McDermott, C. D. Monteiro, G. W. Gundersen, A. Ma’ayan, Enrichr: a comprehensive gene set enrichment analysis web server 2016 update. Nucleic acids research. 44, W90–7 (2016).

69. M. R. Corces, J. D. Buenrostro, B. Wu, P. G. Greenside, S. M. Chan, J. L. Koenig, M. P. Snyder, J. K. Pritchard, A. Kundaje, W. J. Greenleaf, R. Majeti, H. Y. Chang, Lineage-specific and single-cell chromatin accessibility charts human hematopoiesis and leukemia evolution. Nat Genet. 48, 1193–1203 (2016).

70. B. Chapuy, M. R. McKeown, C. Y. Lin, S. Monti, M. G. M. Roemer, J. Qi, P. B. Rahl, H. H. Sun, K. T. Yeda, J. G. Doench, E. Reichert, A. L. Kung, S. J. Rodig, R. A. Young, M. A. Shipp, J. E. Bradner, Discovery and characterization of super-enhancer-associated dependencies in diffuse large B cell lymphoma. Cancer Cell.24, 777–790 (2013).

71. S. G. Landt, G. K. Marinov, A. Kundaje, P. Kheradpour, F. Pauli, S. Batzoglou, B. E. Bernstein, P. Bickel, J. B. Brown, P. Cayting, Y. Chen, G. DeSalvo, C. Epstein, K. I. Fisher-Aylor, G. Euskirchen, M. Gerstein, J. Gertz, A. J. Hartemink, M. M. Hoffman, V. R. Iyer, Y. L. Jung, S. Karmakar, M. Kellis, P. V. Kharchenko, Q. Li, T. Liu, X. S. Liu, L. Ma, A. Milosavljevic, R. M. Myers, P. J. Park, M. J. Pazin, M. D. Perry, D. Raha, T. E. Reddy, J. Rozowsky, N. Shoresh, A. Sidow, M. Slattery, J. A. Stamatoyannopoulos, M. Y. Tolstorukov, K. P. White, S. Xi, P. J. Farnham, J. D. Lieb, B. J. Wold, M. Snyder, ChIP-seq guidelines and practices of the ENCODE and modENCODE consortia. Genome Res. 22, 1813–1831 (2012).

72. B. Egan, C.-C. Yuan, M. L. Craske, P. Labhart, G. D. Guler, D. Arnott, T. M. Maile, J. Busby, C. Henry, T. K. Kelly, C. A. Tindell, S. Jhunjhunwala, F. Zhao, C. Hatton, B. M. Bryant, M. Classon, P. Trojer, An Alternative Approach to ChIP-Seq Normalization Enables Detection of Genome-Wide Changes in Histone H3 Lysine 27 Trimethylation upon EZH2 Inhibition. PloS one. 11, e0166438 (2016).

73. S. Mehta, A. Buyanbat, S. Orkin, B. Nabet, High-efficiency knock-in of degradable tags (dTAG) at endogenous loci in cell lines. Methods Enzymol (2022), doi:10.1016/bs.mie.2022.08.045.

74. J. Navarrete-Perea, Q. Yu, S. P. Gygi, J. A. Paulo, Streamlined Tandem Mass Tag (SL-TMT) Protocol: An Efficient Strategy for Quantitative (Phospho)proteome Profiling Using Tandem Mass Tag-Synchronous Precursor Selection-MS3. J Proteome Res. 17, 2226–2236 (2018).

75. J. Li, Z. Cai, R. D. Bomgarden, I. Pike, K. Kuhn, J. C. Rogers, T. M. Roberts, S. P. Gygi, J. A. Paulo, TMTpro-18plex: The Expanded and Complete Set of TMTpro Reagents for Sample Multiplexing. J Proteome Res. 20, 2964–2972 (2021).

76. D. K. Schweppe, J. K. Eng, Q. Yu, D. Bailey, R. Rad, J. Navarrete-Perea, E. L. Huttlin, B. K. Erickson, J. A. Paulo, S. P. Gygi, Full-Featured, Real-Time Database Searching Platform Enables Fast and Accurate Multiplexed Quantitative Proteomics. J Proteome Res. 19, 2026–2034 (2020).

77. R. Rad, J. Li, J. Mintseris, J. O’Connell, S. P. Gygi, D. K. Schweppe, Improved Monoisotopic Mass Estimation for Deeper Proteome Coverage. J Proteome Res. 20, 591–598 (2021).

78. B. M. Gassaway, J. Li, R. Rad, J. Mintseris, K. Mohler, T. Levy, M. Aguiar, S. A. Beausoleil, J. A. Paulo, J. Rinehart, E. L. Huttlin, S. P. Gygi, A multi-purpose, regenerable, proteome-scale, human phosphoserine resource for phosphoproteomics. Nat Methods. 19, 1371–1375 (2022).

