## Supplementary Figures for "Leukemia core transcriptional circuitry is a sparsely interconnected hierarchy stabilized by incoherent feed-forward loops"

### Supplementary Figure 1

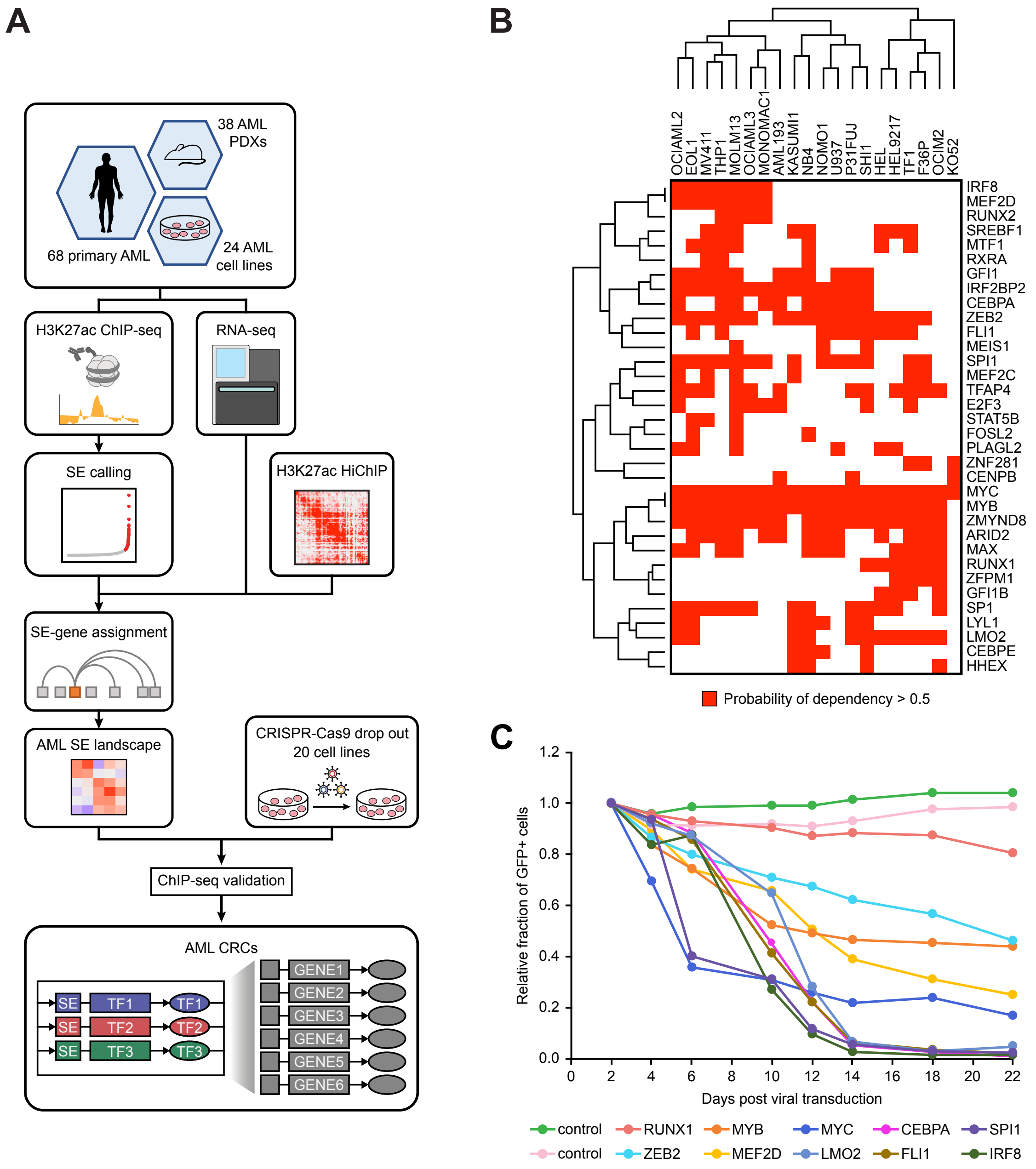

### Supplementary Figure 2

**A**

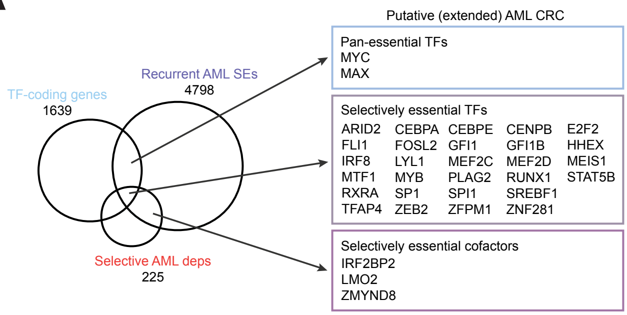


**B**

| CR TF | MV411 dropout score | Probablitiy of dependency in MV411 | Superenhancer in MV411 | Average AML dropout score | Enriched AML dependency | Top 2245 SEs in AML/77% dependency recall | TF (Lambert *et al.*) | ChIP-seq | RNA-seq after k/o in MV411 | dTAG model | Which criteria not met | Reason for inclusion |
|---|---|---|---|---|---|---|---|---|---|---|---|---|
| MYC | -1.9355 | 1 | Y | -2.1153 | N | Y | Y | Y | Y | N | Classified as a common essential gene. | AML oncogene. Strong dependency enrichment in AML despite classification as a common essential. One of the top AML superenhancers |
| MYB | -1.4678 | 1 | Y | -1.3166 | Y | Y | Y | Y | Y | Y | | |
| IRF8 | -0.984 | 0.95 | Y | -0.2734 | Y | Y | Y | Y | Y | Y | | |
| SPI1 | -0.9666 | 0.94 | Y | -0.8617 | Y | Y | Y | Y | Y | Y | | |
| TFAP4 | -0.9586 | 0.94 | N | -0.5679 | Y | Y | Y | Y | N | N | | |
| ARID2 | -0.9466 | 0.94 | Y | -0.5934 | Y | Y | Y | N | N | N | | |
| ZEB2 | -0.9135 | 0.93 | Y | -0.8919 | Y | Y | Y | Y | Y | N | | |
| ZMYND8 | -0.8749 | 0.92 | Y | -0.8435 | Y | Y | N | Y | Y | N | Not classified as a TF by Lambert et al; acts as a transcriptional corepresssor; no evidence of sequence-specific DNA binding | Strong AML dependency; one of the top AML SEs |
| GFI1 | -0.8635 | 0.92 | Y | -0.4162 | Y | Y | Y | Y | Y | Y | | |
| STAT5B | -0.8203 | 0.9 | N | -0.3804 | Y | Y | Y | Y | N | N | | |
| SREBF1 | -0.8202 | 0.9 | Y | -0.5842 | Y | Y | Y | Y | N | N | | |
| MEF2D | -0.7837 | 0.88 | Y | -0.392 | Y | Y | Y | Y | Y | Y | | |
| SPI1 | -0.7638 | 0.86 | N | -0.5498 | Y | Y | Y | Y | Y | N | | |
| IRF2BP2 | -0.7218 | 0.83 | Y | -0.7084 | Y | Y | N | Y | Y | Y | Not classified as a TF by Lambert et al; acts as a transcriptional corepresssor; no evidence of sequence-specific DNA binding | Strong AML dependency; one of the top AML SEs |
| MTF1 | -0.6026 | 0.69 | N | -0.3934 | Y | N | Y | N | N | N | | |
| RXRA | -0.5299 | 0.58 | Y | -0.2658 | Y | Y | Y | Y | N | N | | |
| MEF2C | -0.4962 | 0.52 | Y | -0.2991 | Y | Y | Y | Y | Y | N | | |
| CEBPA | -0.4748 | 0.49 | Y | -0.6419 | Y | Y | Y | Y | N | N | | |
| FOSL2 | -0.4277 | 0.4 | Y | -0.2716 | Y | Y | Y | Y | N | N | | |
| MAX | -0.4247 | 0.39 | N | -0.5456 | N | Y | Y | Y | Y | N | Common essential gene, not a preferential AML dependency | Forms a heterodimer with MYC; strong dependency and SE |
| MEIS1 | -0.4143 | 0.38 | Y | -0.2146 | Y | Y | Y | Y | Y | N | | |
| FLI1 | -0.3815 | 0.32 | Y | -0.4868 | Y | Y | Y | Y | Y | N | | |
| PLAGL2 | -0.3803 | 0.32 | N | -0.423 | Y | Y | Y | Y | N | N | | |
| E2F3 | -0.3061 | 0.2 | Y | -0.5525 | Y | Y | Y | Y | N | N | | |
| RUNX1 | -0.2808 | 0.17 | Y | -0.4783 | Y | Y | Y | Y | Y | Y | | |
| LYL1 | -0.2597 | 0.14 | Y | -0.4766 | Y | Y | Y | Y | Y | N | | |
| RUNX2 | -0.2505 | 0.13 | Y | -0.3184 | Y | Y | Y | Y | Y | Y | | |
| CEBPE | -0.228 | 0.11 | N | -0.4682 | Y | N | Y | N | N | N | | |
| ZNF281 | -0.1853 | 0.07 | N | -0.2791 | Y | Y | Y | Y | N | N | | |
| HHEX | -0.1486 | 0.05 | N | -0.3016 | Y | Y | Y | N | N | N | | |
| GFI1B | -0.1443 | 0.05 | N | -0.2929 | Y | N | Y | N | N | N | | |
| CENPB | -0.1062 | 0.03 | N | -0.2181 | Y | Y | Y | N | N | N | | |
| ZFPM1 | -0.0528 | 0.02 | Y | -0.2008 | Y | Y | Y | N | N | N | | |
| LMO2 | -0.0324 | 0.01 | Y | -0.5689 | Y | Y | N | N | N | N | Not classified as a TF by Lambert et al; acts as a scaffolding cofactor for TF complexes; no evidence of sequence-specific DNA binding | Transcriptional regulator; leukemia oncogene; strong dependency and SE |

### Supplementary Figure 3

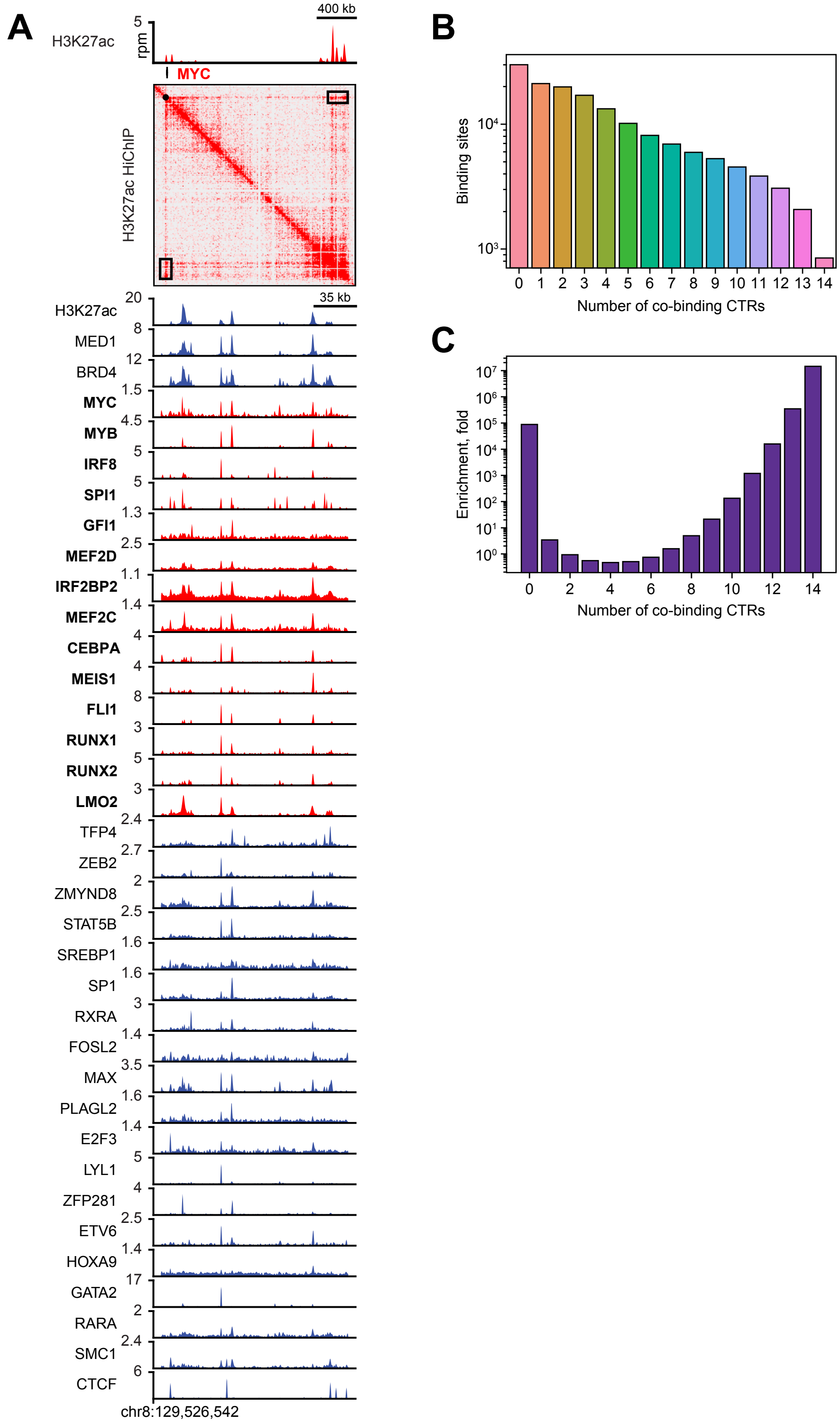

### Supplementary Figure 4

A

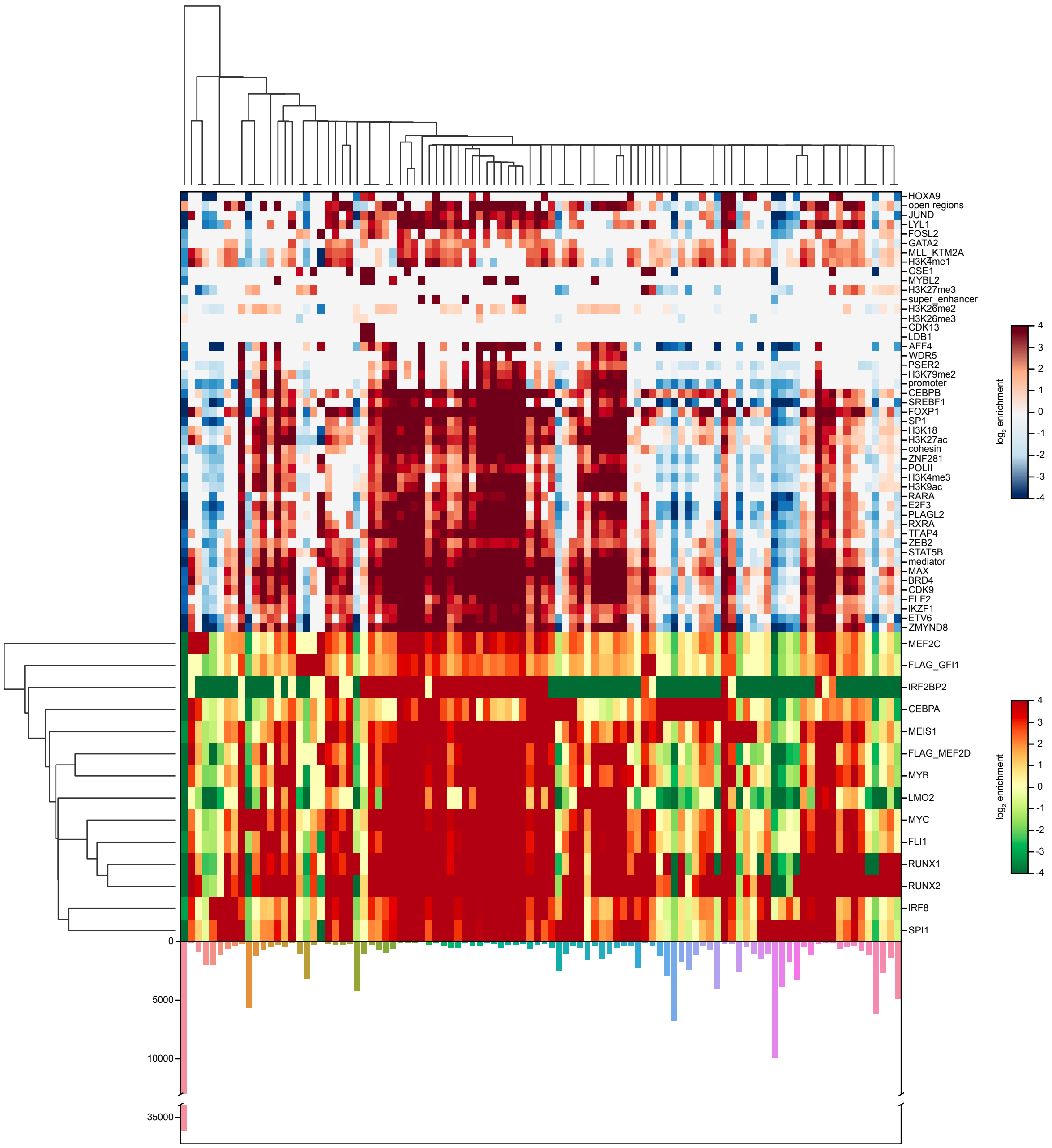

### Supplementary Figure 5

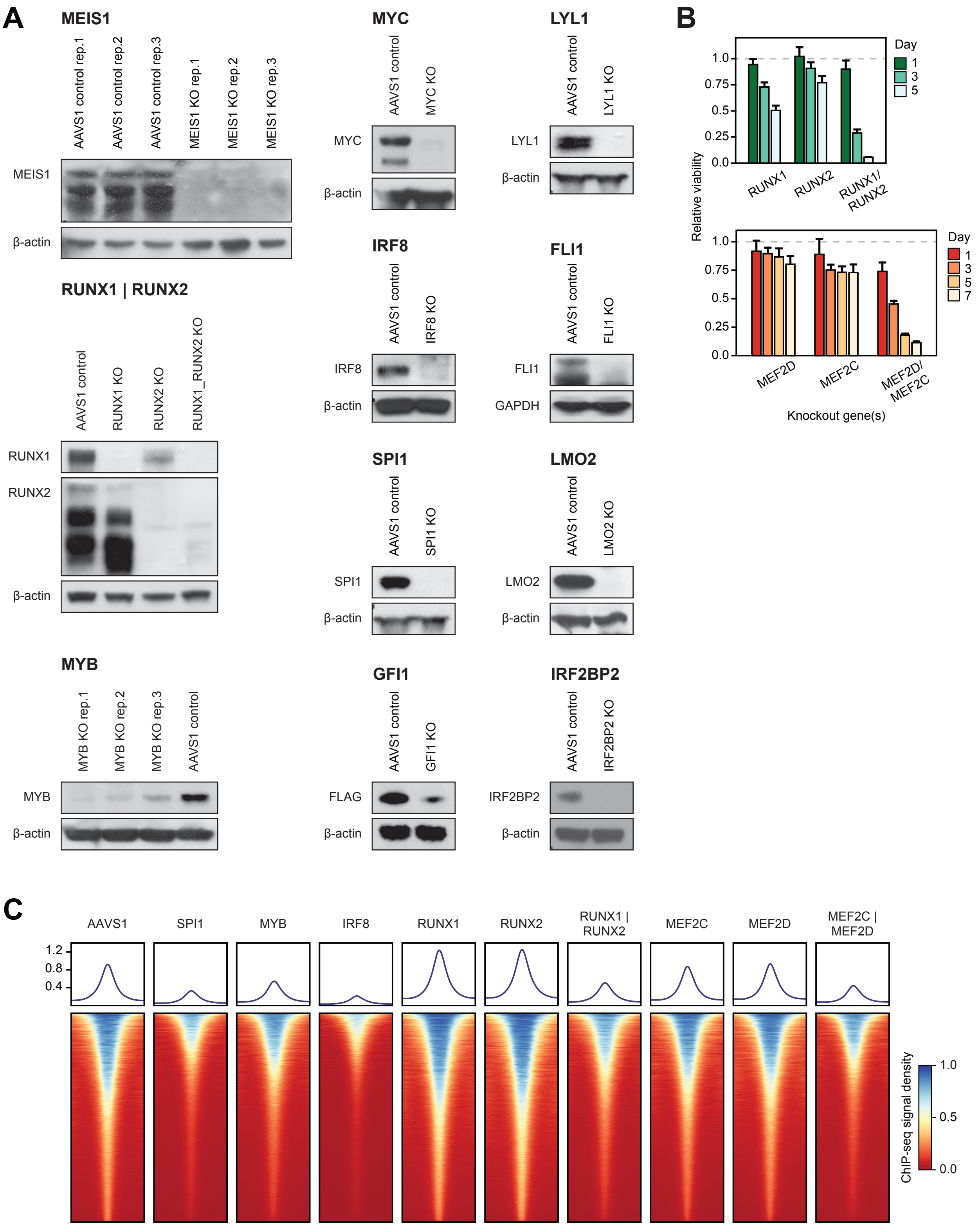

### Supplementary Figure 6

A

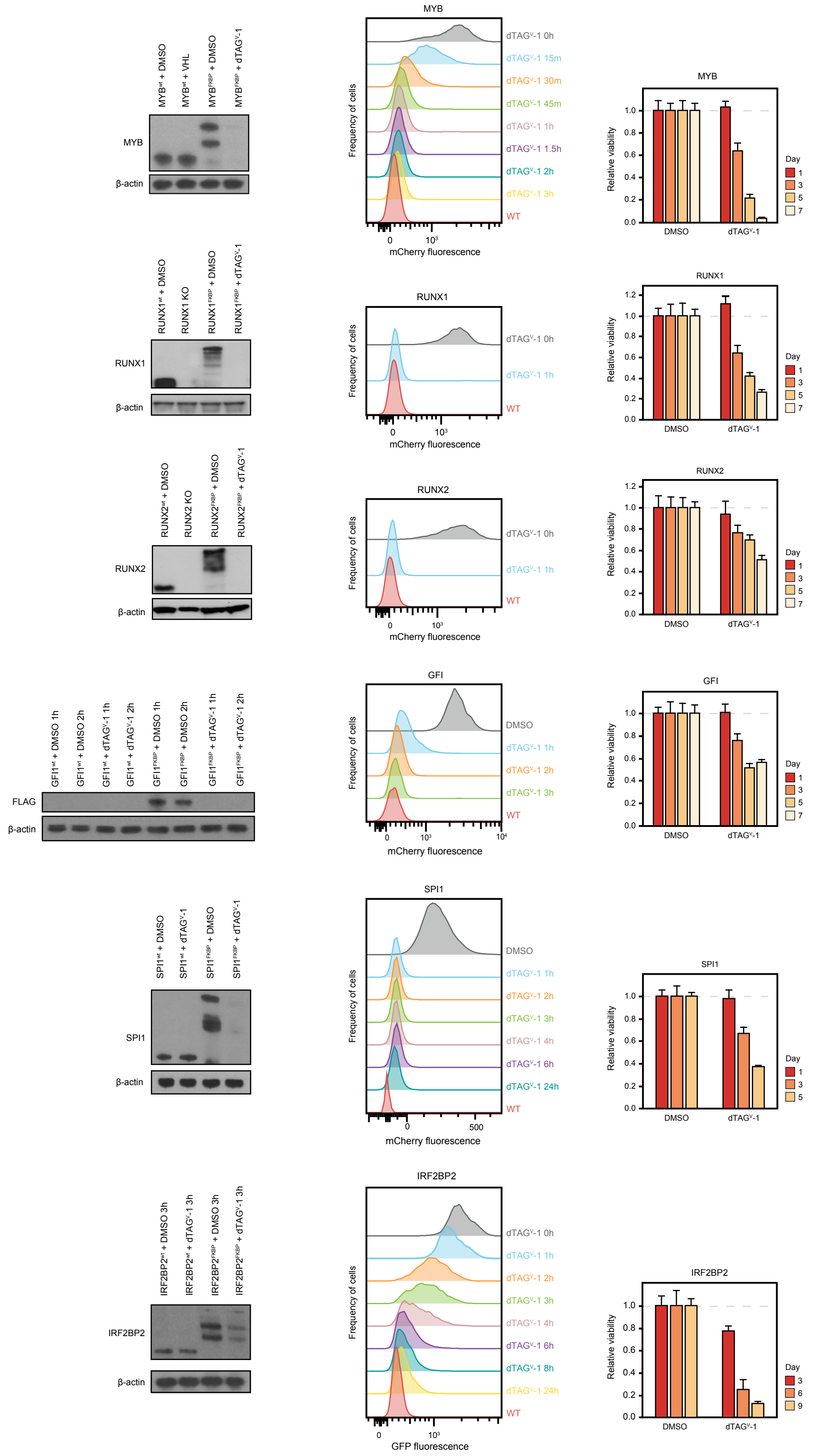

### Supplementary Figure 7

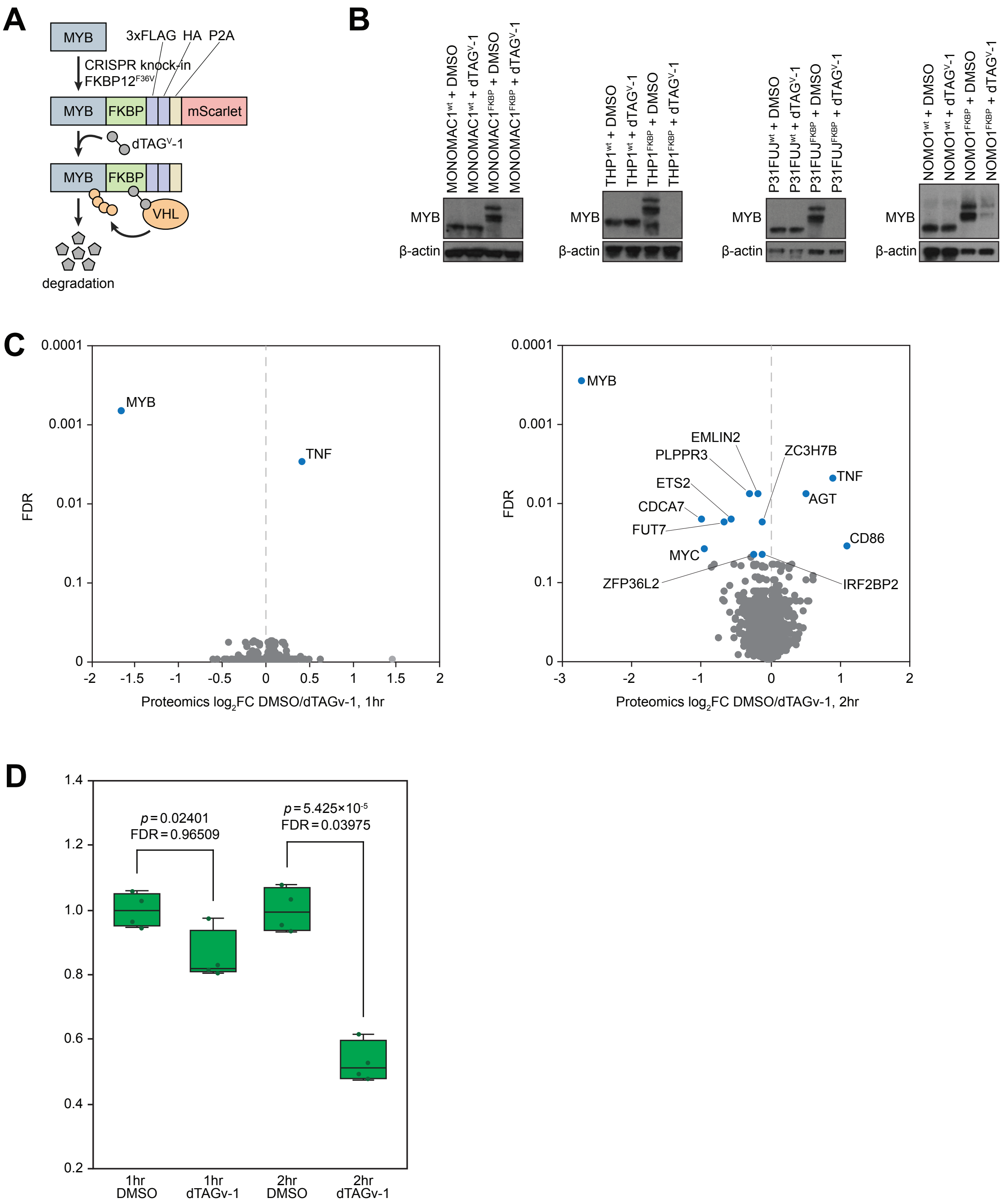

### Supplementary Figure 8

**A**

| | IRF2BP2 | GFI1 | RUNX1 | RUNX2 | SPI1 | IRF8 | MEF2D | MYB_1h | MYB_2h | MYB_4h | MYB_8h | MYB_12h | MYB_24h | MYB_48h | MYB_MONOMAC1 | MYB_NOMO1 | MYB_P31FUJ | MYB_THP1 | JQ1 | THZ1 | THZ1 + JQ1 |
|---|---|---|---|---|---|---|---|---|---|---|---|---|---|---|---|---|---|---|---|---|---|
| IRF2BP2 | 1.00 | 0.12 | 0.27 | 0.12 | 0.28 | 0.18 | 0.60 | 0.14 | 0.09 | 0.08 | 0.07 | 0.08 | 0.05 | 0.04 | 0.12 | 0.10 | 0.12 | 0.14 | 0.07 | 0.13 | 0.07 |
| GFI1 | 0.08 | 1.00 | 0.18 | 0.15 | 0.31 | 0.20 | 0.20 | 0.15 | 0.09 | 0.05 | 0.05 | 0.05 | 0.04 | 0.03 | 0.16 | 0.11 | 0.10 | 0.16 | 0.04 | 0.07 | 0.05 |
| RUNX1 | 0.10 | 0.09 | 1.00 | 0.20 | 0.18 | 0.16 | 0.16 | 0.10 | 0.05 | 0.03 | 0.03 | 0.03 | 0.02 | 0.01 | 0.10 | 0.05 | 0.06 | 0.10 | 0.03 | 0.07 | 0.03 |
| RUNX2 | 0.10 | 0.18 | 0.46 | 1.00 | 0.22 | 0.20 | 0.20 | 0.12 | 0.07 | 0.06 | 0.05 | 0.06 | 0.03 | 0.03 | 0.15 | 0.10 | 0.11 | 0.11 | 0.05 | 0.09 | 0.05 |
| SPI1 | 0.09 | 0.15 | 0.17 | 0.09 | 1.00 | 0.38 | 0.38 | 0.12 | 0.05 | 0.03 | 0.03 | 0.04 | 0.02 | 0.01 | 0.10 | 0.07 | 0.09 | 0.12 | 0.03 | 0.06 | 0.03 |
| IRF8 | 0.03 | 0.05 | 0.07 | 0.04 | 0.18 | 1.00 | 0.00 | 0.05 | 0.02 | 0.01 | 0.02 | 0.02 | 0.01 | 0.01 | 0.05 | 0.03 | 0.03 | 0.07 | 0.01 | 0.02 | 0.01 |
| MEF2D | 0.03 | 0.01 | 0.07 | 0.04 | 0.03 | 0.00 | 1.00 | 0.02 | 0.01 | 0.01 | 0.00 | 0.01 | 0.00 | 0.00 | 0.01 | 0.01 | 0.01 | 0.02 | 0.01 | 0.01 | 0.00 |
| MYB_1h | 0.20 | 0.31 | 0.38 | 0.21 | 0.54 | 0.42 | 0.67 | 1.00 | 0.31 | 0.17 | 0.15 | 0.18 | 0.09 | 0.07 | 0.43 | 0.30 | 0.30 | 0.52 | 0.09 | 0.17 | 0.10 |
| MYB_2h | 0.31 | 0.50 | 0.48 | 0.32 | 0.58 | 0.52 | 0.47 | 0.79 | 1.00 | 0.35 | 0.36 | 0.35 | 0.21 | 0.17 | 0.56 | 0.47 | 0.42 | 0.59 | 0.20 | 0.34 | 0.21 |
| MYB_4h | 0.52 | 0.46 | 0.53 | 0.44 | 0.60 | 0.54 | 0.67 | 0.74 | 0.59 | 1.00 | 0.50 | 0.59 | 0.30 | 0.25 | 0.50 | 0.37 | 0.35 | 0.51 | 0.27 | 0.44 | 0.30 |
| MYB_8h | 0.46 | 0.47 | 0.54 | 0.39 | 0.58 | 0.64 | 0.60 | 0.69 | 0.64 | 0.53 | 1.00 | 0.73 | 0.37 | 0.28 | 0.58 | 0.46 | 0.43 | 0.62 | 0.31 | 0.51 | 0.34 |
| MYB_12h | 0.36 | 0.34 | 0.42 | 0.34 | 0.49 | 0.54 | 0.60 | 0.53 | 0.42 | 0.42 | 0.49 | 1.00 | 0.28 | 0.19 | 0.41 | 0.30 | 0.28 | 0.42 | 0.23 | 0.36 | 0.24 |
| MYB_24h | 0.57 | 0.62 | 0.62 | 0.46 | 0.65 | 0.76 | 0.87 | 0.78 | 0.70 | 0.60 | 0.70 | 0.78 | 1.00 | 0.60 | 0.62 | 0.52 | 0.51 | 0.69 | 0.46 | 0.59 | 0.52 |
| MYB_48h | 0.66 | 0.69 | 0.65 | 0.59 | 0.65 | 0.72 | 0.73 | 0.81 | 0.75 | 0.66 | 0.69 | 0.72 | 0.80 | 1.00 | 0.69 | 0.58 | 0.58 | 0.72 | 0.55 | 0.62 | 0.62 |
| MYB_MONOMAC1 | 0.15 | 0.29 | 0.37 | 0.24 | 0.39 | 0.40 | 0.33 | 0.38 | 0.20 | 0.10 | 0.11 | 0.12 | 0.06 | 0.05 | 1.00 | 0.33 | 0.31 | 0.54 | 0.09 | 0.14 | 0.08 |
| MYB_NOMO1 | 0.20 | 0.32 | 0.31 | 0.25 | 0.44 | 0.44 | 0.47 | 0.43 | 0.27 | 0.13 | 0.15 | 0.14 | 0.09 | 0.08 | 0.54 | 1.00 | 0.52 | 0.61 | 0.09 | 0.14 | 0.10 |
| MYB_P31FUJ | 0.18 | 0.21 | 0.26 | 0.20 | 0.38 | 0.30 | 0.40 | 0.30 | 0.17 | 0.08 | 0.10 | 0.09 | 0.06 | 0.05 | 0.35 | 0.36 | 1.00 | 0.49 | 0.09 | 0.11 | 0.09 |
| MYB_THP1 | 0.08 | 0.13 | 0.16 | 0.08 | 0.21 | 0.24 | 0.20 | 0.21 | 0.09 | 0.05 | 0.06 | 0.06 | 0.03 | 0.03 | 0.25 | 0.17 | 0.20 | 1.00 | 0.04 | 0.07 | 0.04 |
| JQ1 | 0.45 | 0.42 | 0.64 | 0.45 | 0.61 | 0.56 | 0.73 | 0.43 | 0.37 | 0.31 | 0.33 | 0.36 | 0.26 | 0.23 | 0.47 | 0.31 | 0.41 | 0.44 | 1.00 | 0.56 | 0.60 |
| THZ1 | 0.23 | 0.19 | 0.37 | 0.20 | 0.34 | 0.18 | 0.40 | 0.21 | 0.17 | 0.13 | 0.14 | 0.16 | 0.09 | 0.07 | 0.21 | 0.12 | 0.14 | 0.21 | 0.15 | 1.00 | 0.17 |
| THZ1 + JQ1 | 0.47 | 0.46 | 0.51 | 0.38 | 0.62 | 0.54 | 0.67 | 0.46 | 0.40 | 0.33 | 0.36 | 0.38 | 0.29 | 0.26 | 0.44 | 0.32 | 0.42 | 0.42 | 0.59 | 0.62 | 1.00 |

**B**

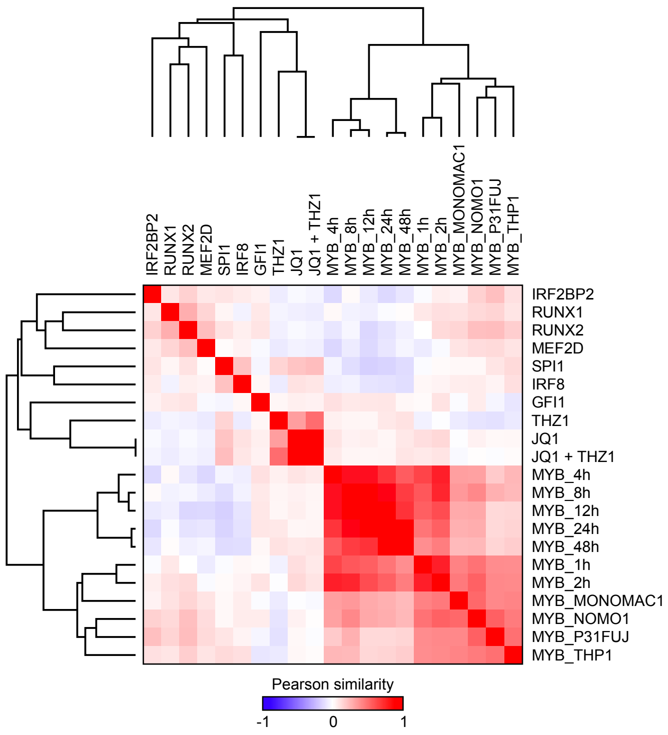

### Supplementary Figure 9

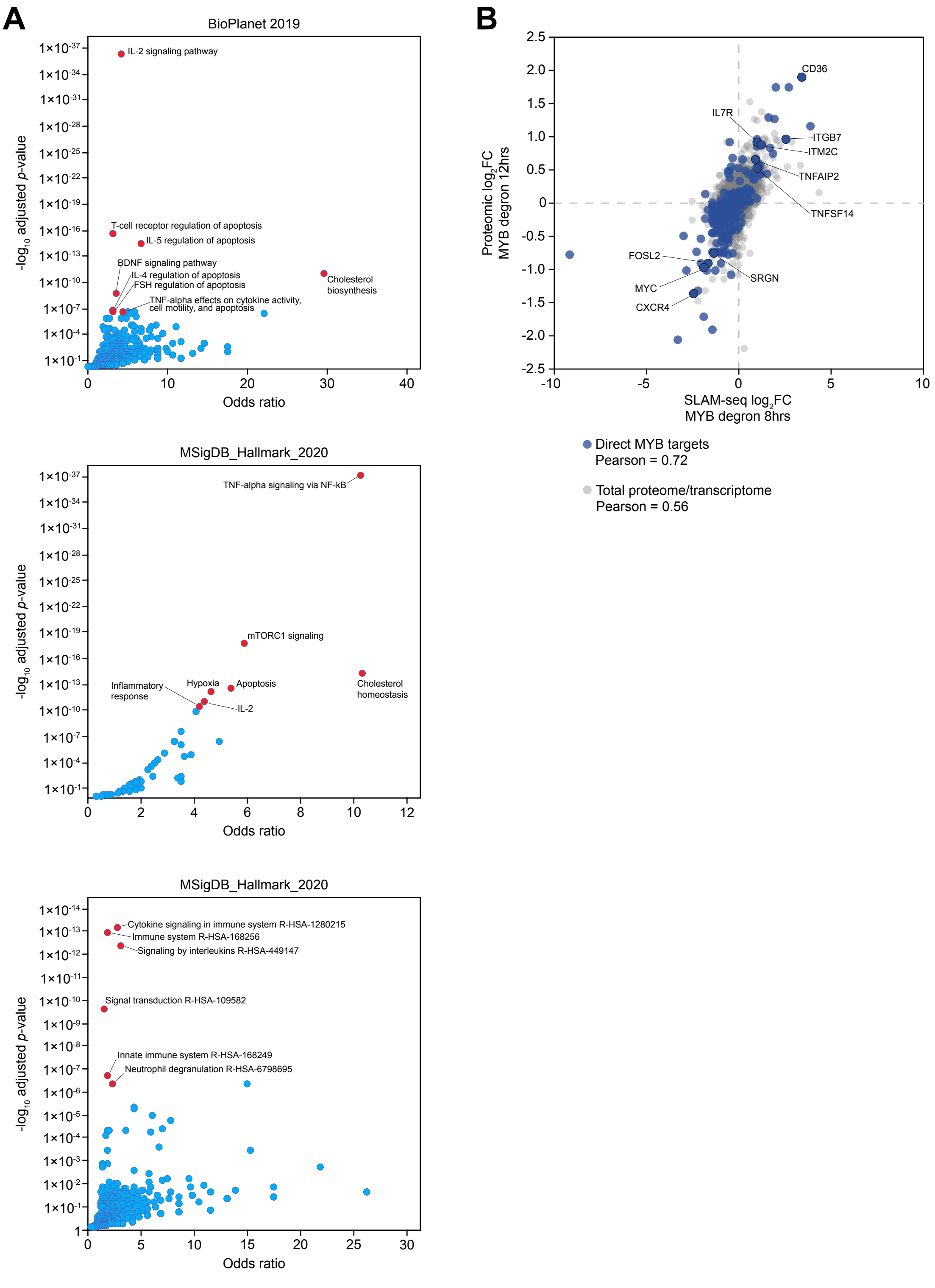

### Supplementary Figure 10

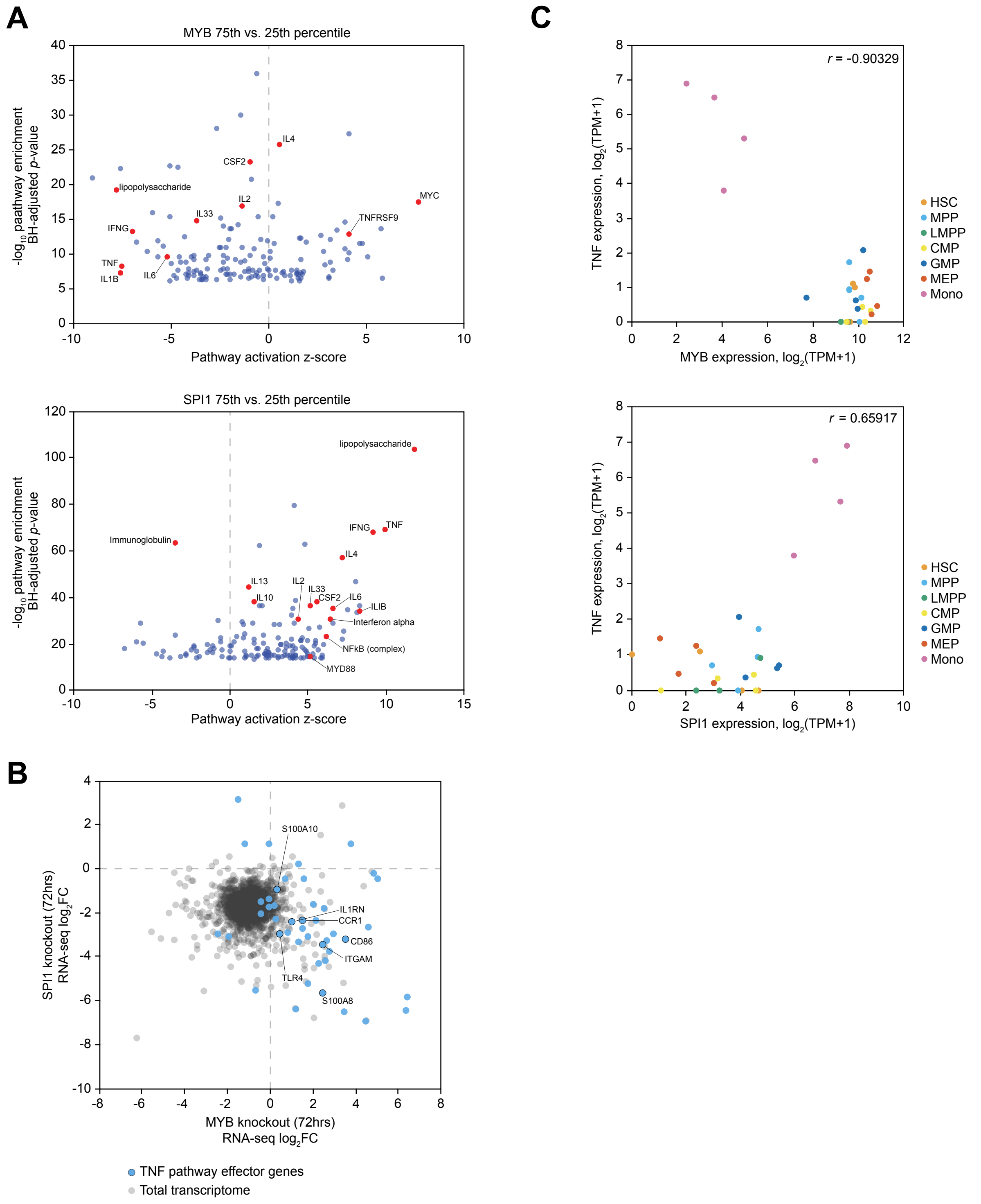
